# Phenotype-dependent subtyping exposes high MYC activity as a targetable dependency in a large subset of human Lung Adenocarcinoma

**DOI:** 10.64898/2026.04.27.721038

**Authors:** Sarah Laing, Isabel Dye, Ryan Byrne, Eva C. Freckmann, Robin Shaw, Shahnawaz Ali, Sophie Heese, Yoana Doncheva, Catherine Ficken, Jennifer Doig, Sudhir B Malla, Saira Ghafoor, Sophie McLaughlin, Colin Nixon, Graeme Clark, Leah Officer-Jones, Emily Hoey, Claire Kennedy-Dietrich, Nicola Brady, Doug Strathdee, Iain A. McNeish, John Le Quesne, Crispin Miller, Karen Blyth, Nigel B. Jamieson, Matthew J. Fuchter, Philip Dunne, Robert Brown, Daniel J. Murphy

## Abstract

C-MYC (MYC) occupies a critical nexus of oncogenic signalling and deregulated expression of MYC is widespread across most human cancer types, suggesting that MYC should be an attractive target for therapeutic intervention. Although 30-40% of human Non-Small Cell lung cancers show low level amplification of *c-MYC* and genetic evidence has shown that *c-Myc* is a key downstream effector of KRas-driven lung tumourigenesis in mouse models, the functional contribution of MYC to human lung cancer remains unclear. We applied a phenotype-based classifier to the TCGA Lung Adenocarcinoma (LuAd) cohort and found that high MYC transcriptional activity identifies a subset of LuAd with significantly reduced survival. Application of the same methodology to a panel of genetically engineered mouse models identified multiple genotypes that give rise to the high MYC activity phenotype, disease positioning such models as reflective of a distinct subset of human LuAd. We show that high MYC activity predicts sensitivity to a small molecule dual-inhibitor of the MYC co-factors, EZH2 and G9A, HKMTi-1-005, and that treatment with HKMTi-1-005 strongly reduced MYC protein expression, induced B cell-mediated immune surveillance and suppressed growth of autochthonous KRas^G12D^-driven lung tumours.

**Statement of significance:** This work establishes the principle of indirectly targeting MYC in LuAd, via inhibition of associated enzymatic cofactors, EZH2 and G9A, and identifies a large subset of aggressive human LuAd with a high MYC activity signature that may benefit from this approach.

## INTRODUCTION

The oncogene *cMYC* (henceforth MYC) is well-established to contribute to the growth, evolution, and resistance to treatment of a broad spectrum of human cancers, both solid and blood-borne (1–3). The MYC coding sequence is rarely mutated, and overexpression is the predominant mode through which MYC contributes to cancer. Overexpression of MYC may arise from copy number gains, chromosomal translocation, increased transcription, translation, or protein stability, and is frequently achieved via oncogenic mutations in upstream regulators, including EGFR, KRAS, APC, PI3K, AKT and NOTCH (4–8). Subtle increases in MYC expression suffice to drive ectopic proliferation of most terminally differentiated adult cell types *in vivo* and have been shown to accelerate tumour development in multiple adult tissues (9–14). In human lung cancer, over 1/3^rd^ of TCGA-annotated Adenocarcinoma and Squamous Cell carcinoma samples harbour low copy number amplification of the *MYC* locus (15), while genetic experiments in mice have consistently demonstrated that endogenously expressed Myc is a critical effector of both activated KRas and Wnt pathway signalling in LuAd (16–18), reflecting multiple routes to MYC involvement in lung cancer.

While MYC participates in additional activities, such as DNA replication and RNA splicing (19, 20), MYC is first and foremost a transcription factor, regulating the expression of genes involved in almost every aspect of cell biology (21). To fulfil this role, MYC must heterodimerise with MAX and bind in a sequence selective manner to DNA. When MYC levels are low, binding is largely limited to high-affinity DNA sites, such as E-box motifs. As MYC levels rise, binding across the genome becomes increasingly promiscuous, broadening the impact of MYC on the transcriptome (22). In normal cells, MYC levels are held in check by the fact that elevated expression of MYC is a potent trigger of canonical apoptosis, thereby eliminating cells that exceed a tolerable threshold of MYC expression (9, 23). As nascent tumours emerge and progress, mutations such as KRAS activation, p53 inactivation, or upregulation of anti-apoptotic BCL2 family proteins, raise the threshold level of MYC required to trigger apoptosis, allowing tumour cells to tolerate higher MYC expression (24). Such mutations may have combinatorial effects in this regard, for instance by modulating expression or function of different members of the BCL2 homology domain 3 (BH3) family of apoptotic regulators, upon which canonical MYC-induced apoptosis largely depends (25–28). In principle, this should allow MYC levels to continue rising without triggering apoptosis as tumours accumulate more mutations and progress to more aggressive disease.

Pharmacological efforts to target MYC have heretofore focussed largely on either disrupting MYC:MAX heterodimer formation or limiting expression of MYC, the latter by transcriptional, translational, or post-translational means (29). For instance, Bromodomain inhibitors can reduce MYC transcription, while Sylvestrol has been shown to block MYC translation through stabilisation of a G-quadruplex structure in the 5’UTR of *MYC* mRNA (30, 31). MYC protein function and stability is regulated by an array of Ubiquitin ligases and Deubiquitylases (32, 33), many of which are amenable to small molecule inhibition, however, the considerable redundancy in the network presents a clear challenge to achieving effective MYC depletion (34). More direct approaches include small molecule inhibitors that prevent MYC dimerization with MAX, such as MYCMI6, MYCMI7 and MYCi975 (35–37), and the peptide inhibitor OMO-103 (aka OMO-MYC), the latter being the only MYC inhibitor to date to show activity in clinical trials (38).

An approach that has garnered less attention is to target the enzymatic activity of co-factors required for MYC to regulate transcription. Alongside its better-known activity as a positive regulator of gene expression, MYC also forms transcription repressing complexes (39), notably with MIZ1, encoded by *ZBTB17*, and a Histone methyltransferase, G9A, encoded by *EHMT2* (40, 41), or alternatively, with EZH2, also a Histone methyltransferase and the enzymatic subunit of Polycomb Repressor Complex 2 (42). Through formation of these complexes MYC represses expression of a broad array of genes, notably including cyclin-dependent kinase inhibitors, such as p21 and p15 (43, 44), and genes involved in immune visibility, such as Type I Interferon regulators and MHC I antigen presentation pathway genes (45). Indeed, transcriptional repression of immune visibility by MYC has been reported across numerous cancer types, including pancreatic, breast, lung and liver cancers (11, 18, 46–49) and is evident within 24hrs of MYC upregulation in MEFs (11), suggesting that immune evasion is a core activity of MYC-mediated transcriptional regulation. In this regards, selective suppression of repressive MYC complexes may be an attractive therapeutic approach, particularly if combined with immune checkpoint blockade, especially given that immune effector cells also require the transactivating functions of MYC for their proliferation (50).

The low level of MYC amplification in lung cancer, combined with the potent pro-tumourigenic activity of even modestly overexpressed MYC, present major obstacles to identifying a lung cancer patient cohort that would benefit from a MYC-targeted therapy. Here we apply a phenotype-driven subtyping approach to stratify patients based on Hallmark Pathways gene expression and identify an aggressive subtype of human LuAd defined by high MYC transcriptional activity. We show that this high MYC activity subtype is readily observable in widely-used mouse models of LuAd and test the strategy of indirectly targeting MYC activity in pre-clinical models of LuAd, using small molecule inhibition of its repressive co-factors, G9A/EZH2.

## RESULTS

### MYC transcriptional activity defines an aggressive subtype of human Lung Adenocarcinoma

Molecular subtyping of lung cancer is currently based primarily on the presence or absence of specific recurring mutations in driver oncogenes, e.g., EGFR, KRAS, BRAF or ALK. A large subset of human lung Adenocarcinoma (LuAd) however lacks obvious driver mutations, limiting rational approaches to treatment for such cases. A recently developed pathway-based classifier approach revealed novel phenotypic subtypes of colorectal cancer, irrespective of mutation status, that may better reflect underlying tumour biology than gene-level analysis (51). We therefore applied pathway-derived subtyping (PDS) methodologies to the TCGA cohort of human LuAd (N = 537). Using the *PDSclassifier* R package, we identified three distinct subtypes (PDS1-PDS3) with high confidence, along with a minor subgroup that failed to meet the prediction threshold for any individual subtype, labelled as “mixed” (Fig. 1A). Gene expression levels for MSigDB Hallmarks signature “MYC Targets V1” is significantly elevated in PDS1 LuAd tumours, whereas “KRAS Signalling Down” is significantly elevated in PDS3 LuAd tumours, compared to the other two classes (Fig. 1B). PDS1 tumours additionally showed the highest proliferative index and replication stress signatures (Fig. S1A). Using the full N=50 Hallmarks from the MSigDB collection (52), we observed strong enrichment of MYC Targets V1/V2, Glycolysis, MTORC1 and cell-cycle-related signalling in PDS1 tumours; EMT and Inflammation-related signalling pathways in PDS2 tumours; and KRAS Signalling Down, Bile Acid and Fatty Acid Metabolism enriched in PDS3 tumours (Fig. 1C). Accordingly, MYC and E2F family transcription factors are highly active in PDS1, whereas FOS, JUN, STAT and IRF family transcription factors are most active in PDS2 tumours (Fig. S1B). Unlike the genotype-independent nature of PDS in colorectal cancer, PDS3 LuAd tumours are significantly enriched for mutations in KRAS (p=0.0009), EGFR (p=0.00725) and STK11 (p=0.0237), whereas PDS1 tumours are enriched for Tp53 mutation (p=7.39e^-13^). PDS2 tumours are notably enriched for wild type STK11 (p=0.008) (Fig. 1C). Using scores generated for MYC targets V1 and MYC targets V2 in these tumours, stratified into tertiles, we observe stepwise increases for the highest expression tertile across stage I-IV tumours (Fig. 1D, E). Using clinical follow-up data for these tumours, we observe a stepwise association between the levels of Hallmark MYC Targets V1, or MYC Targets V2, and patient outcome, with significantly lower survival associated with the highest tertile in each instance (Fig. 1F). Higher MYC activity thus correlates significantly with more aggressive LuAd.

**Figure 1.**
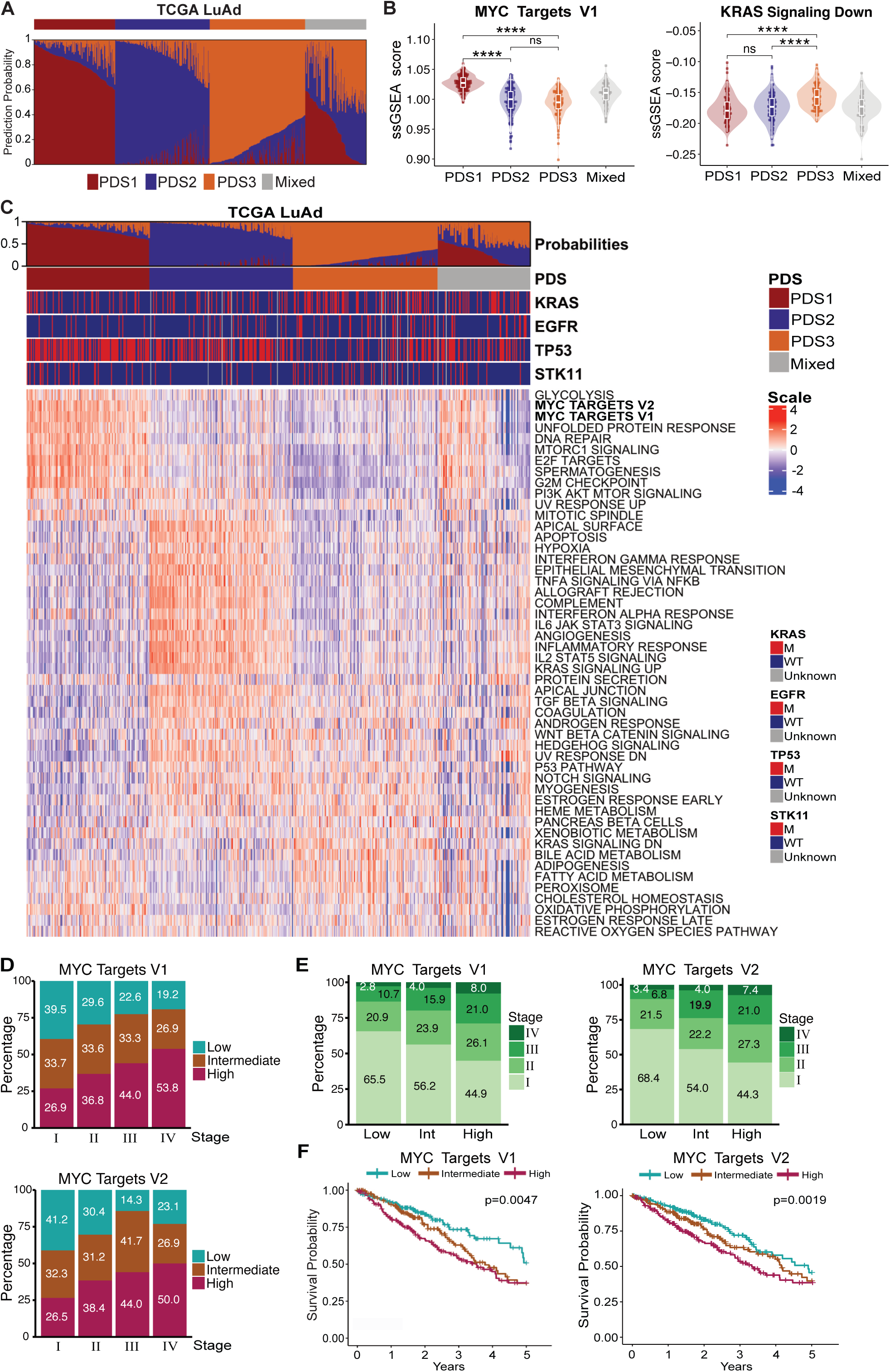
**A)** Barplot illustrating the Pathway-Derived Subtype (PDS) prediction probabilities for each of the TCGA lung adenocarcinoma (LuAd) samples (n=537). Each vertical bar represents a single tumour sample, with colours indicating the predicted probabilities of the given sample being a member of each of the three subtypes (red = PDS1, blue = PDS2, orange = PDS3). The top annotation bar indicates the assigned PDS, with samples displaying a probability > 0.6 of belonging to a given PDS being assigned to that PDS. Samples failing to reach this assignment threshold (i.e., probability < 0.6 for all three subtypes) were classified as Mixed (grey). For clarity, samples are grouped by assigned subtype and then within each assigned subtype ordered by descending prediction probability. **B)** ssGSEA scores for MYC Targets V1 (left) and KRAS Signaling Dn (right) Hallmarks according to PDS subtype in TCGA LuAd. Each dot represents 1 tumour sample. Boxplots indicate interquartile range (25th–75th percentiles) with median represented by the central horizontal line. Whiskers extend to values within 1.5 times the interquartile range of the upper and lower quartiles. Observations beyond the end of the whiskers are plotted individually. Mann-Whitney test values indicated. **C)** Heatmap showing ssGSEA scores for all 50 Hallmark gene sets in the TCGA LuAd samples, with samples arranged according to PDS. Annotation bars above heatmap indicating mutational status of KRAS, EGFR, TP53 and STK11 in each sample (M = Mutant, WT = Wild type). **D)** (Top) Proportion of samples with low, intermediate and high MYC Targets V1 ssGSEA scores within each LuAd stage: Stage I (n=294), Stage II (n=125), Stage III (n=84) and Stage IV (n=26). Stage information was absent for 8 samples that were excluded from this analysis. (Bottom) Proportion of samples with low, intermediate and high MYC Targets V2 ssGSEA scores within each LuAd stage. Samples were divided into tertiles (low, intermediate and high) based on ssGSEA scores. (Left) Stage distribution within the low, intermediate, and high MYC Targets V1 groups. (Right) LuAd stage distribution within the low (N=177), intermediate (N=176), and high (N=176) MYC Targets V2 groups. **E)** Five-year overall survival according to MYC Targets V1 (left) and MYC Targets V2 (right) low (N=177), intermediate (N=178) and high (N=173) groups. Survival data was absent for 9 samples which were excluded from this analysis. Log rank test p-value shown. For all panels, **** denotes p<0.0001, *** = p<0.001, ** = p<0.01, * = p<0.05, and ns = p>0.05.

### Disease-positioning of genetically engineered mouse models of LuAd

Strictly contingent upon exposure to CRE recombinase, the lox-stop-lox-KRas^G12D^ allele expresses constitutively active KRas from its endogenous locus and has served as a mainstay of *in vivo* LuAd research for over 2 decades (53). In a typical experiment, tumours are initiated sporadically in the lungs of adult mice by intranasal installation of replication-incapacitated viral vectors expressing CRE recombinase. Following viral delivery of CRE to the lungs, activation of KRas^G12D^ alone gives rise to multiple independent hyperplastic lesions, a small subset of which progress to Adenocarcinoma over a protracted period (53). The allele is therefore widely used in combination with additional CRE-activated oncogenic (e.g. *Rosa26^DM.lsl-MYC^*) or CRE-inactivated tumour suppressor (e.g. *Tp53^fl/fl^*, *STK11^fl/fl^*) cooperating alleles to accelerate tumour development (10, 54–56). The inclusion of a conditional allele for expression of human APOBEC3B confers immune visibility to these models (57, 58). We assembled a panel of such combinatorial allelic mice, induced sporadic lung tumours by intranasal installation of Adeno-SPC-CRE, and harvested tumours for analysis when mice reached humane clinical end-points. Following histological examination of lungs, we generated a tissue microarray (TMA) reflecting the diversity of tumour morphological phenotypes (Fig. S2A, B). TMAs were stained with fluorescent labelled pan-Cytokeratin antibody and selected for whole transcriptome RNA-SEQ analysis using the Nanostring GeoMX workflow. The diffuse nature of lung tissue allowed for ready distinction between tumours and adjacent normal tissue. While selecting regions of interest we noted a third morphologically distinct phenotype representing focal areas of hyperplastic lung tissue, present in all genotypic cohorts, which we included in our analysis (Fig. S2C). Regions of interest (ROIs) were collected (Table S1) without further segmentation, sequenced and subject to comparative whole transcriptome analysis. Principal component analysis showed hyperplastic and normal tissue clustering together, away from the spread of tumour phenotypes (Fig. S3A). Notably, tumours from different genotypes did not segregate into distinct clusters but instead all nucleated around a core cluster surrounded by a scattering of individual tumours of various genotypes with no obvious pattern, even following removal of the normal and hyperplastic samples (Figure S3A, B). We applied PDS subtyping based on Hallmark Pathways to the dataset, as per the human analysis above. Roughly 60% of murine tumours could be designated into PDS1-3 subtypes, as per human LuAd, with a stringent probability threshold of 0.6. Consistent with the human PDS1 data showing enrichment of Tp53 mutations and MYC activity, PDS1 tumours arose largely from KM^2^, KP and KPA mice (Figure 2A). Interestingly, tumours from KM mice with only 1 copy of *Rosa26^DM.lsl-MYC^* fell largely into PDS3 or the mixed phenotype subtype, suggesting that a threshold level of MYC deregulation is required to drive the PDS1 “high MYC” phenotype (Fig. 2A and S3C). Tumours with loss of p53 exhibited the greatest phenotypic plasticity and were present in all 3 PDS subtypes along with the mixed phenotype group. Curiously, the PDS2 “inflammatory” subtype was enriched for tumours with deletion of STK11 (KL & KLA), at odds with the significant enrichment of wild-type STK11 in human PDS2 LuAd.

**Figure 2.**
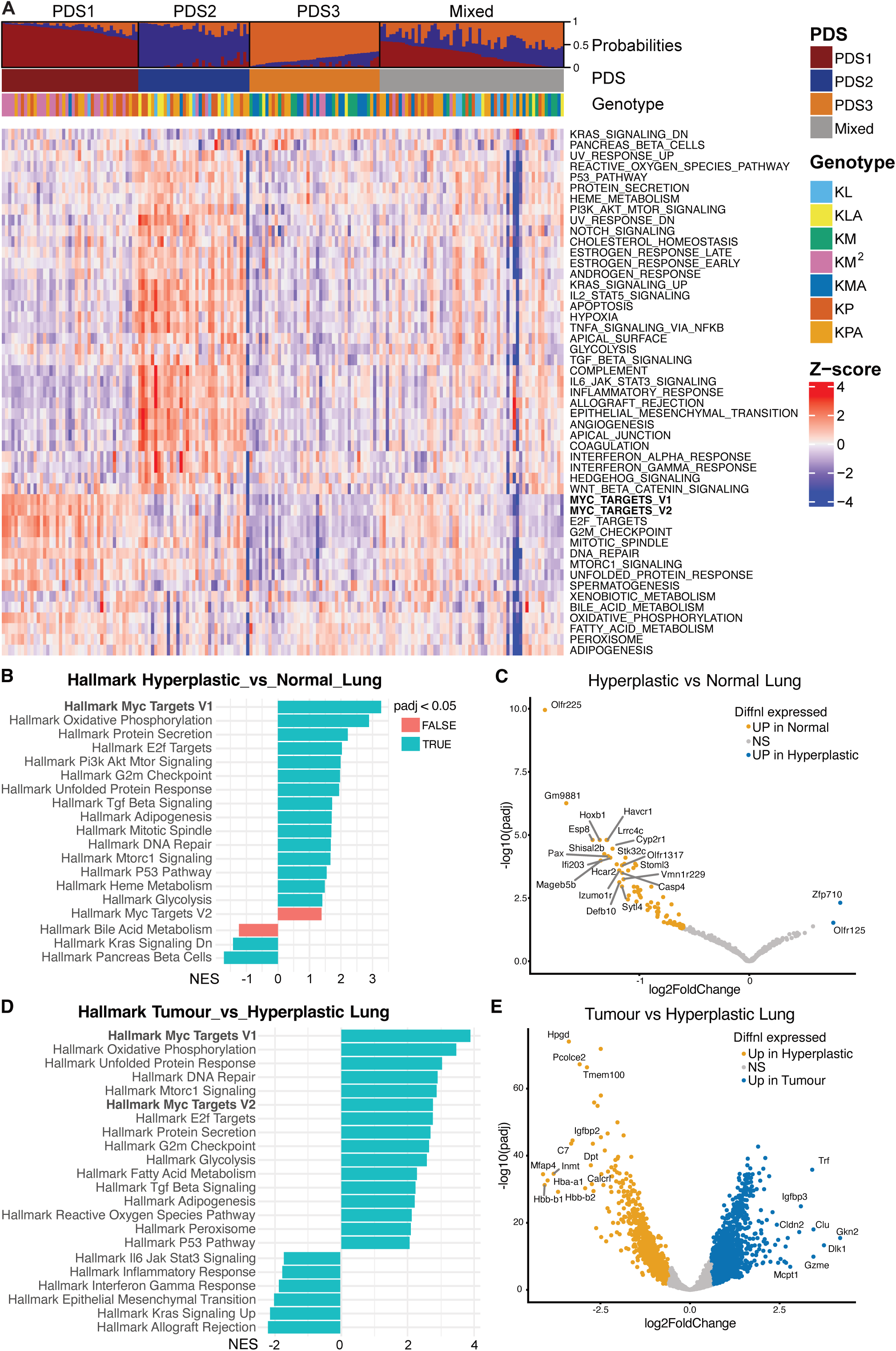
**A)** Heatmap showing ssGSEA scores for all 50 Hallmark gene sets in tumour samples from TMAs of LuAd mouse models, with samples stratified to PDS subtypes, as described in Fig 1C. Annotation bars above heatmap indicating the genotype of the mouse tumour sample. **B)** Differential expression analysis showing ssGSEA scores for Hallmark pathways that are differentially expressed in mouse samples of all genotypes between hyperplastic lung tissue and adjacent normal lung tissue. FDR<0.05 indicated by turquoise bars. **C)** Volcano plot indicating genes that are differentially expressed in mouse samples of all genotypes between hyperplastic lung tissue and adjacent normal lung tissue. Genes showing significantly differential expression (2x or more, FDR<0.05) highlighted in orange and blue. **D)** Differential expression analysis showing ssGSEA scores for Hallmark pathways that are differentially expressed in mouse samples of all genotypes between lung tumour tissue and hyperplastic lung tissue. FDR<0.05 indicated by turquoise bars. **E)** Volcano plot indicating genes that are differentially expressed in mouse samples of all genotypes between lung tumour tissue and hyperplastic lung tissue. Genes showing significantly differential expression (2x or more, FDR<0.05) highlighted in orange and blue.

We also investigated the transcriptomic differences between hyperplastic versus normal ROIs, along with hyperplastic versus tumour ROIs. Strikingly, “Hallmark MYC Targets V1” and Hallmark pathways that are typically enriched following acute MYC upregulation (e.g., Hallmark E2F Targets; MTORC1 Signalling; G2M Checkpoint) were amongst the most highly enriched pathways in the more aggressive phenotype for both comparisons (Fig. 2B-E) and remained so upon re-analysis following removal of all *Rosa26^DM.lsl-MYC^* expressing tumour samples (Fig. S3D). Expression of endogenous *c-Myc* was not different between normal and hyperplastic tissue but was significantly elevated in KP, KPA, KL & KLA tumours, relative to normal and hyperplastic tissue (Fig S3E). Note that tumours induced in mice carrying the *Rosa26^DM.lsl-MYC^* allele were omitted from this analysis to avoid the confounding effect of exogenous MYC expression on endogenous *Myc* levels (59). We conclude from these data that stepwise increases in MYC activity define key transitions from the very outset of lung tumour initiation, continuing to increase as tumours progress to aggressive late-stage cancer.

### MYC levels predict sensitivity to G9A/EZH2 inhibition

The Cancer Dependency Map (https://depmap.org/portal) Achilles project (60) indicates a high requirement for MYC expression in non-small cell lung cancer cell lines, with CRISPR-mediated *MYC* deletion yielding a median Chronos score of −1.99 (interquartile range −2.24 to −1.67), well below the dependency threshold of −0.5. Of 98 NSCLC cell lines in the Achilles database, only 4 fail to show dependency for *cMYC* (Fig. 3A). We selected H358 cells as representative of median MYC dependency and A549 as representative of high MYC dependency and attempted direct pharmacological inhibition of MYC using a previously published small molecule, MYCi975, in these cell lines. Despite previous evidence of *in vivo* efficacy against other cancer types (12, 35), MYCi975 achieved only modest inhibition of lung tumour cell proliferation *in vitro* at the highest concentration tested, with slightly greater sensitivity noted in A549s compared with H358 cells, consistent with DepMap data (Fig. 3B). We therefore considered indirect approaches to targeting MYC, via pharmacological inhibition of MYC co-factors.

**Figure 3.**
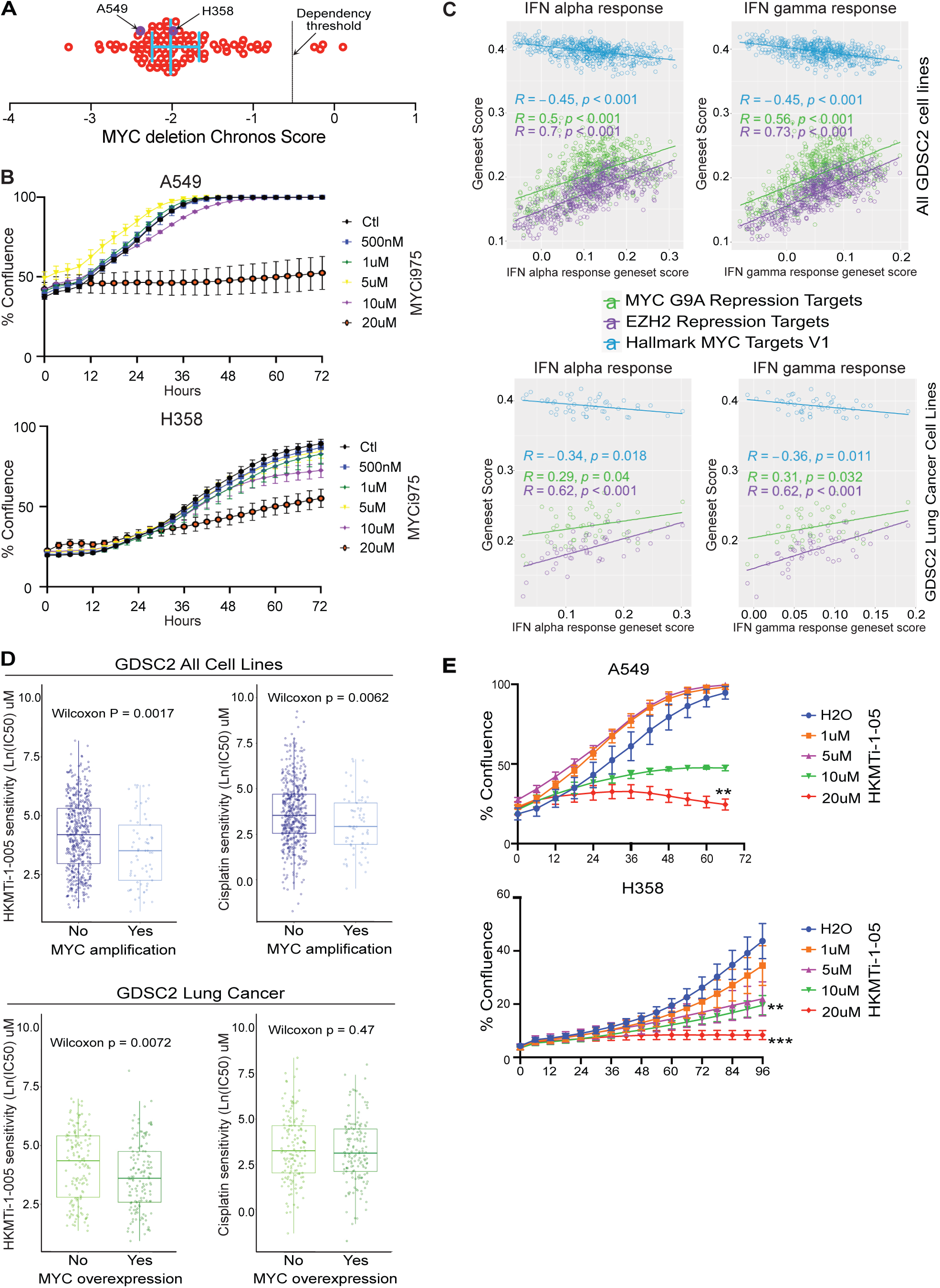
**A)** Chronos scores from the DepMap Achilles project for 98 Non-Small Cell Lung Cancer cell lines following CRISPR-mediated deletion of cMYC. Error bars indicate median Chronos score and interquartile range, relative to dependency threshold (0.5). **B)** Confluence of A549 and H358 cells treated with 500nm, 1µM, 5 µM, 10 µM & 20 µM MYCi975 or DMSO vehicle measured over time by Incucyte. Mean values ± SEM of 9 technical replicates (3 image fields x 3 replicate wells) shown. Data are representative of 4 independent experiments. **C)** Spearman’s correlation between Interferon Alpha Response (left panels), or Interferon Gamma Response (right panels) genesets (x axis) and genesets reflecting targets of MYC-G9a repression (in green), EZH2 repression (purple), and hallmark MYC activation (blue). Each circle represents geneset scores in 1 cell line across all GDSC2 solid tumour cell lines (upper panels; N = 565) or GDSC2 lung cancer cell lines (lower panels; N = 49). **D)** Difference in sensitivity to HKMTi-1-005 or cisplatin between cMYC-amplified (N = 65) and non-amplified (N = 502) solid tumour cell lines (upper panels) and between highest and lowest quartiles for MYC expression (lower panels; N = 141 each). Wilcoxon test. **E)** Confluence of A549 and H358 cells treated with 1 µM, 5 µM, 10 µM and 20 µM HKMTi-1-005 or H_2_O vehicle measured over time by Incucyte. Mean values ± SEM of three independent experiments shown. Significant difference versus vehicle controls determined using one-way ANOVA with post-hoc Tukey test for multiple comparisons.

The Histone methyltransferases G9A and EZH2 form separate repressive complexes with MYC (41, 42) and SMIs of both are available. Having previously shown that MYC complexes directly repress expression of Interferon pathway transcription factors, STAT1, STAT2, IRF7 and IRF9 in multiple settings (11), we asked if there was existing evidence of regulation of the IFN pathway by G9A or EZH2. *In silico* analysis of the Genomics of Drug Sensitivity in Cancer 2 (GDSC2) pan-cancer collection of 565 solid tumour cell lines (61) revealed strong positive correlations between “Interferon Alpha Response”/ “Interferon Gamma Response”/ “Innate Immune Response” gene sets and “MYC G9A Repression Targets”, suggesting that MYC-dependent repression of Interferon signalling is a general feature of solid tumours. Similarly strong positive correlations between “Interferon Alpha Response”/ “Interferon Gamma Response”/ “Innate Immune Response” gene sets and “EZH2 Repression Targets” were also observed, along with strong negative correlations with the canonical (positively regulated) “Hallmark MYC Targets V1” gene set (Fig. 3C top panels, & Fig. S4A left panel). Limiting this analysis to GDSC2 lung cancer cell lines mirrored these correlations, albeit to a lesser but nevertheless significant degree (Fig. 3C bottom panels, & Fig. S4A right panel). Notably, an experimental dual inhibitor of G9A and EZH2, HKMTi-1-005 was previously shown to induce Interferon signalling and limit tumour growth in a mouse model of Ovarian cancer (62). Using the GDSC2 drug sensitivity data we asked if MYC levels predict sensitivity to HKMTi-1-005. Across all tumour types, MYC amplified cell lines (N=65) exhibit significantly greater drug sensitivity than non-amplified lines (N=502), comparable to their higher sensitivity to Cisplatin (Fig. 3D upper panels). MYC amplified lines moreover exhibit significantly elevated sensitivity to individual inhibitors of G9A (UNC0638) or EZH2 (GSK353; Fig. S4B left panels). Given that MYC is frequently overexpressed in the absence of gene amplification, we compared drug sensitivity across cell lines in the highest and lowest quartiles for MYC expression (N=141 for each). Cell lines in the highest quartile for MYC expression exhibit significantly elevated sensitivity to HKMTi-1-005, or to the individual G9A inhibitor UNC0638, but not to Cisplatin or the individual EZH2 inhibitor, GSK353 (Fig 3D lower panels, & Fig. S4B right panels), compared with cell lines in the lowest quartile. To verify these data in lung cancer cells lines, we performed a titration of all 3 drugs, HKMTi-1-005, GSK353 and UNC0638, and measured growth inhibition of A549 and H358 cells by Incucyte longitudinal imaging. HKMTi-01005 marginally outperformed the other 2 compounds across both cell lines, achieving growth inhibition that was comparable to the combined effect of GSK353 and UNC0638 (Fig. 3E & Fig. S4C-E).

### HKMTi-1-005 induces Interferon signalling in murine lung tumours

The KM mouse model of Lung Adenocarcinoma combines conditional expression of constitutively active KRas^G12D^ with that of human cMYC, driven conditionally from the Rosa26 locus (*Rosa26^DM.lsl-MYC^*) at levels that combine with endogenously expressed *c-Myc* to approximate *MYC* trisomy (11, 12). The KM model exhibits accelerated progression to LuAd, compared with KRas^G12D^ expression alone, and has previously exposed a clinically relevant signature of early LuAd progression (63, 64). Inclusion of a conditional allele expressing APOBEC3B (A3B) from the other copy of the Rosa26 locus (*Rosa26^DS.lsl-A3B^*) yields a transiently immune visible model of autochthonous LuAd (abbreviated “KMA”), without otherwise altering the tumour phenotype (bioRxiv 2023.07.24.550274). Replacing the A3B allele with a second copy of *Rosa26^DM.lsl-MYC^* (KM^2^), again results in accelerated progression to Adenocarcinoma, compared with the KM and KMA models, but at the expense of immune visibility afforded by the A3B allele (10 & bioRxiv 2023.07.24.550274); see schematic, Fig. 4A). We therefore chose the KMA model to ask if HKMTi-1-005 treatment can reverse MYC mediated suppression of Interferon signalling and enhance immune visibility of autochthonous lung tumours. Immune visibility of the KMA model is characterised by pronounced, albeit transient, infiltration of tumours by CD8 and CD4 T cells, accompanied by B cell rich tertiary lymphoid structures adjacent to tumour tissue, evident at 8 weeks following tumour initiation, but completely suppressed by 12 weeks post induction (bioRxiv 2023.07.24.550274). We therefore treated KMA mice with HKMTi-1-005 for 5 consecutive days, immediately before reaching the 12-week post induction timepoint, and harvested lungs for analysis by IHC and bulk RNA-SEQ. Immunohistochemistry for Ki67 showed significantly reduced tumour cell proliferation following HKMTi-1-005 treatment (Fig. 4B & C). We performed whole transcriptome analysis of bulk tumour tissue harvested from the same mice. Metacore GeneGO pathway analysis revealed Interferon Signalling, Complement, and MHC I Antigen presentation pathways to be the top-most upregulated pathways following HKMTi-1-005 treatment of lung tumours (Fig 4D). At the level of individual genes, we measured significantly increased expression of Interferon regulators Stat2, Irf7 and Irf9, along with multiple Interferon induced genes, including the Oligoadenylate Synthase (Oas) gene family (Fig. 4E-G), and MHC I antigen presentation genes H2-T24 and B2m (Fig. S5A, S5B), consistent with multiple reports of MYC-dependent repression of these pathways across cancer types (11, 18, 46, 47, 49). These effects were accompanied by sharply reduced expression of multiple Cyclins, CDK1 and CDK2 (Fig. 4H). Although T cell related gene expression was unchanged following HKMTi-1-005 treatment (not shown), we found a pronounced increase in B cell related gene expression, including significantly increased expression of IghG1 and IghG2c, indicative of mature B cells (Fig. S5C). Accordingly, IHC showed increased tumour infiltration by IgG-expressing B cells following HKMTi-1-005 treatment, although B cell staining in tumour-associated tertiary lymphoid structures were comparable between the 2 groups (Fig. S5D). HKMTi-1-005 treatment thus increased Interferon signalling and B cell, but not T cell, tumour engagement while also reducing tumour cell proliferation.

**Figure 4.**
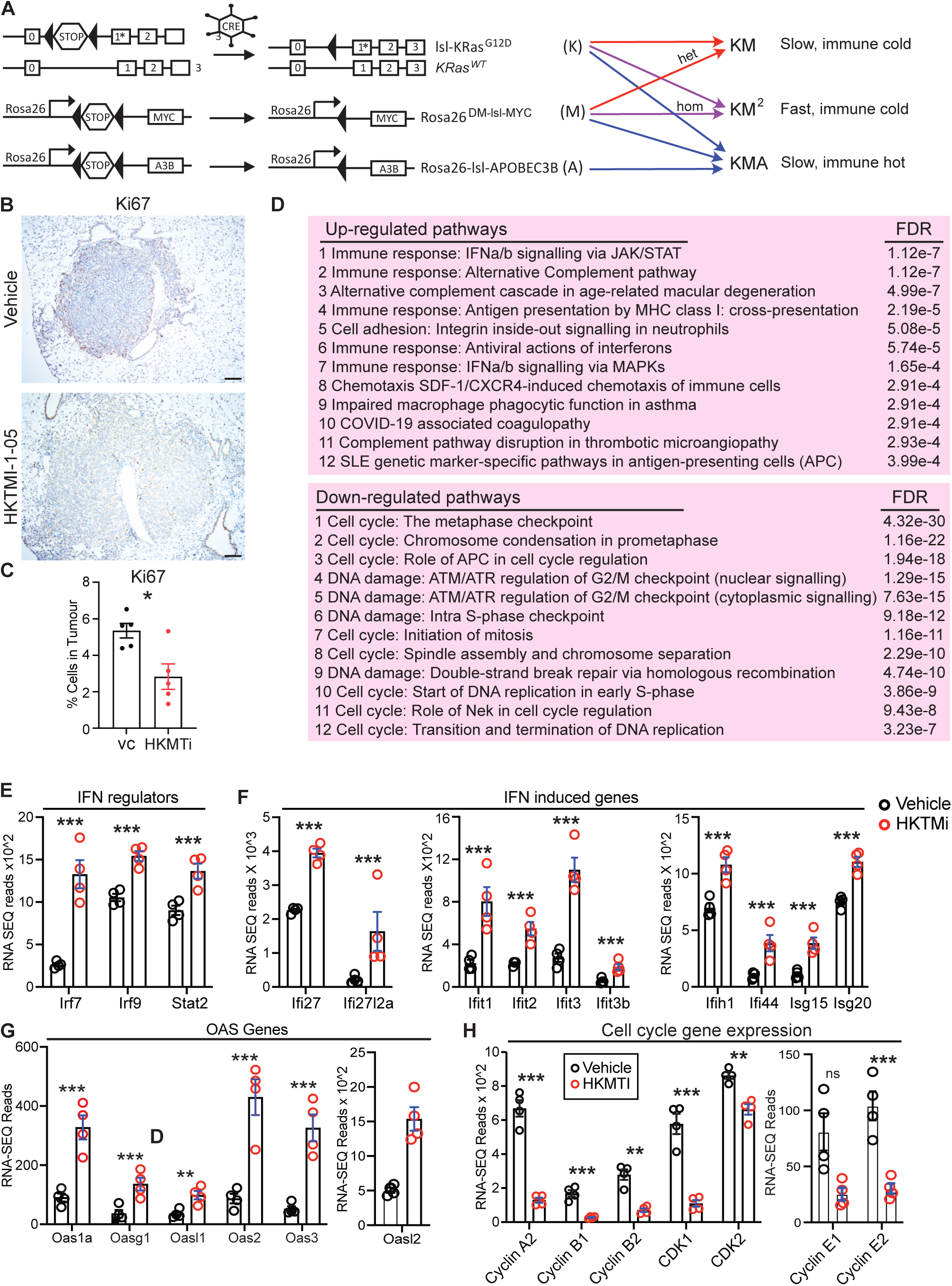
**A)** Schematic of alleles used in Figures 4 & 5. **B)** Representative images of Ki67 staining of KMA tumours following treatment with 40mg/kg HKMTi-1-005 (N=5) or vehicle (N=5) twice daily for 5 days prior to cull at 12-weeks post allele induction (day 78 to day 83). Scale bar 100 µM. **C)** HALO quantification of percentage of Ki67 positive cells in tumours (N = 5 mice in each cohort). Each data point represents the average score of all tumours in a section from an individual mouse. Error bars denote Mean ± SEM. Significance calculated using a T-Test. **D)** The top 12 most significantly up-regulated and down-regulated pathways in tumour bearing KMA mice following treatment with 40mg/kg HKMTi-1-005 or vehicle twice daily for 5 days prior to cull at 12-weeks post allele induction. Significantly modulated pathways were identified using MetaCore GeneGo analysis. **E-H)** Normalised RNA-Seq reads of IFN regulator (E), IFN-induced (F, G) and selected Cell Cycle-related (H) genes from tumour bearing lungs from KMA mice treated with vehicle (N = 4) or 40mg/kg HKMTI-1-005 (N = 4) for 5 days prior to cull at 12-weeks post allele induction. Mean and SEM shown. P values are adjusted for multiple comparisons (Benjamini-Hochberg Test). For all panels, * denotes p<0.05; ** p<0.01; *** p<0.001. NS = not significant.

### Sustained HKMTi-1-005 treatment suppresses growth of MYC-driven LuAd

To evaluate the anti-tumour potential of HKMTi-1-005 we chose the KM^2^ model which, based on our *in silico* analysis above, we reasoned may show increased drug sensitivity compared with the KMA model due to higher MYC expression and a correspondingly higher proliferation index (9, 63). Titration of HKMTi-1-005 showed progressively greater inhibition of tumour cell proliferation in the lungs of KM^2^ mice, with correspondingly reduced lung tumour burden, following 4 weeks treatment with increasing doses of HKMTi-1-005 (Fig. 5A, B). Surprisingly, IHC for MYC revealed sharply reduced protein expression following HKMTi-1-005 treatment (Fig, 5C). Using a daily dose of 35mg/kg HKMTi-1-005, we performed a survival benefit study, treating KM^2^ mice for 4 weeks followed by cessation of treatment, and either harvested lungs for analysis immediately following treatment cessation, or followed mice until humane end points were reached (Fig. 5D). 4-week treatment with HKMTi-1-005 significantly reduced lung tumour burden and tumour cell proliferation (Fig. 5E-H) and moreover significantly prolonged survival of KM^2^ mice, with 7 of 9 treated mice outlasting mice dosed with vehicle (Fig. 5I). We conclude from these experiments that pharmacological inhibition of G9A/EZH2 may be a promising approach for treatment of LuAd with high MYC activity.

**Figure 5.**
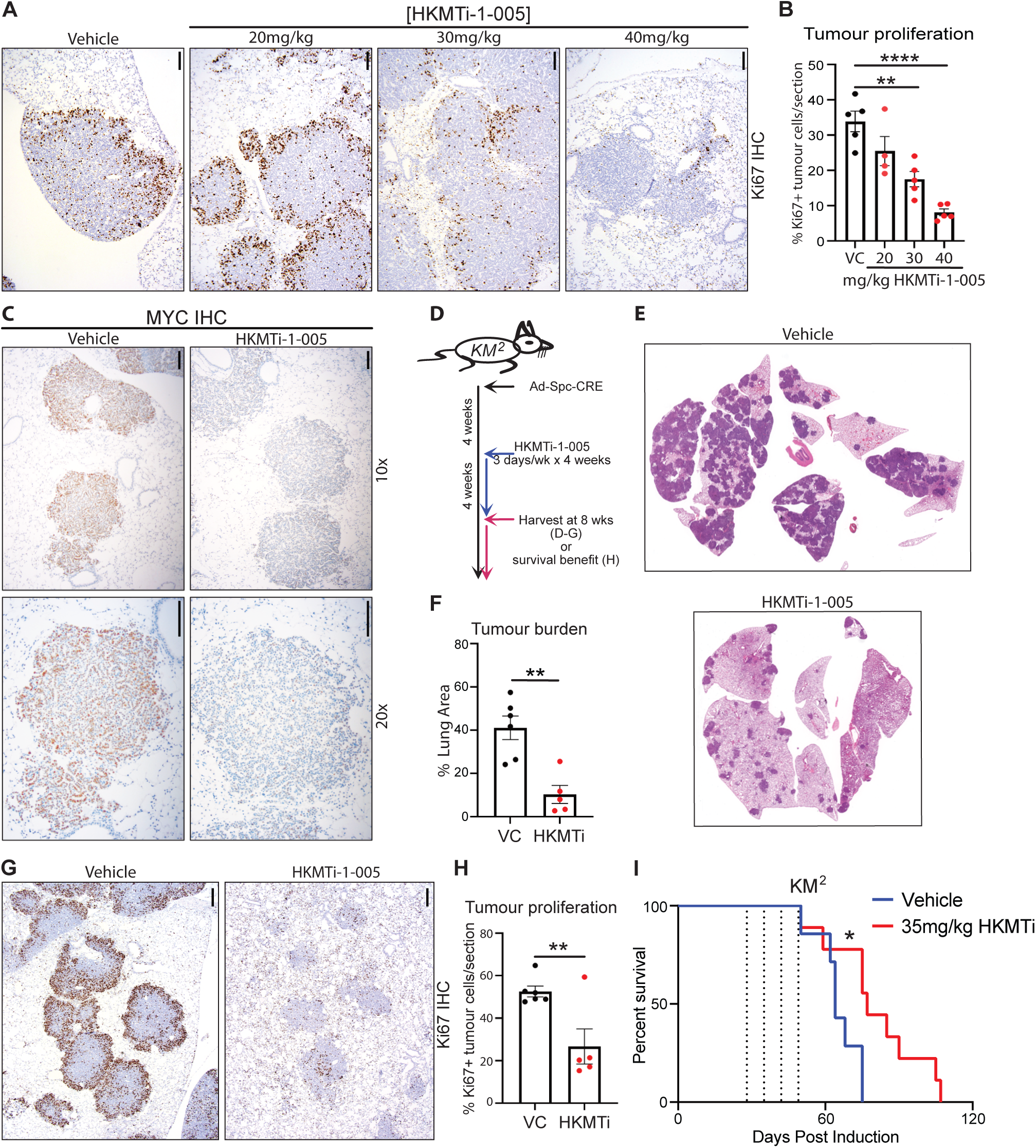
**A)** Representative images of Ki67 staining of KM^2^ tumours following treatment with 20mg/kg (N=4), 30mg/kg (N=4) and 40mg/kg (N=5) HKMTi-1-005 or vehicle (N=5) twice daily for 5 days prior to cull at 8-weeks post allele induction (day 50 to day 55). Scale bars = 100 µM. **B)** HALO quantification of percentage of Ki67+ cells in tumours treated as per (A). Each data point represents the average score of all tumours in a section from an individual mouse. Error bars denote Mean ± SEM. 1 way ANOVA and post-hoc Tukey Test. **C)** Representative images of c-MYC staining of KM^2^ tumours following treatment with 40mg/kg HKMTi-1-005 (N=5) or vehicle (N=5) twice daily for 5 days prior to cull at 8-weeks post allele induction (day 50 to day 55). Scale bars = 100µM. **D)** Schematic showing treatment strategies for KM^2^ mice with HKMTi-1-005. **E)** Representative images of H&E stained lung sections from KM^2^ mice following 4 weeks of 3 days/week treatment with 35mg/kg HKMTi-1-005 (N=5) or vehicle (N=6) and culled on day 56 post allele induction. **F)** Quantification of overall lung tumour burden performed using HALO image analysis software calculated as a percentage of total lung area from KM^2^ mice treated as per (E). Error bars denote Mean ± SEM. Significance calculated using a T-Test. **G)** Representative images of Ki67 staining of KM^2^ tumours from mice treated as per (E, F). Scale bars = 100 µM. N = 6 vehicle and N = 5 HKMTi-1-005. **H)** HALO quantification of percentage of Ki67+ cells in KM^2^ tumours. Each data point represents the average score of all tumours in a section from an individual mouse. Error bars denote Mean ± SEM. Significance calculated using a T-Test. **I)** Overall survival of KM^2^ mice treated with 35mg/kg HKMTi-1-005 (N = 8) or vehicle (N = 10) for 4 weeks and maintained until humane end point following treatment cessation. Treatment interval indicated by the vertical dashed lines. Mantel-Cox log rank test. For all panels, * denotes p<0.05; ** p<0.01; *** p<0.001. NS = not significant.

## DISCUSSION

Cancers of the lung remain the number one cause of death by cancer worldwide. Distinct mutation patterns are prevalent in different geographic regions (e.g., KRAS in western society; EGFR in east Asian communities) and in smokers versus non-smokers (65). Lung cancer in never-smokers comprises an already substantial and growing proportion of cases, while in western society, changes in smoking habits are likely to drive corresponding changes in the mutation spectra present in cases here. Treatment options are currently based largely on the availability of agents targeting specific mutations in EGFR, ALK1, BRAF, and most recently, KRAS, with immunotherapy applied often non-specifically to the remaining half of LuAd cases that lack an actionable “driver” mutation (66, 67). Resistance to targeted agents emerges rapidly while the proportion of cases with durable responses to immunotherapy remains disappointingly low and unpredictable. Improving upon current outcome projections will require development of new strategies for both patient stratification and rationally guided treatments.

Here we use the MSigDB Hallmark Pathways to define 3 phenotypic subtypes of LuAd, with a high MYC activity subtype emerging as significantly more aggressive than LuAd with lower MYC activity. Although certain recurring mutations are enriched in each subtype (e.g., Tp53 in PDS1 and PDS2; KRAS, EGFR and STK11 in PDS3), no exclusive relationship between mutation and subtype is evident. This is borne out by our application of the same methodology to a panel of genetically engineered murine lung tumours, whose gene expression and pathway activity profiles segregate more clearly without reference to genotype than if first stratified by genotype, despite having far less time to evolve than human counterparts. This has two major implications. At the scientific level, GE mouse model users commonly compare phenotypes according to mouse genotype, inferring that differences in phenotype arise as an immediate and direct consequence of genotypic differences. Our data challenge this practice, clearly showing that, while significant differences in gene expression and pathway activity can certainly be identified between genotypes, 1) multiple different genotypes can give rise to convergent phenotypes and 2) any given genotype can result in a spectrum of phenotypes. This latter point is especially true for tumours engineered through loss of Tp53, which exhibit extreme phenotypic plasticity. At the clinical level, treatment selections based entirely on the mutation status of a single gene are thus unlikely to result in uniform patient benefit. The addition of a phenotype-based classifier to complement driver mutation data may improve patient stratification leading to better treatment selection.

Our data now expose a heretofore unappreciated contribution of increasing MYC transcriptional activity to all stages of LuAd tumour progression, from early neoplastic emergence to progressive enrichment of MYC activity in late stage, more aggressive LuAd. The prominence of MYC pathway signatures differentiating between normal and hyperplastic tissue, and again between hyperplastic tissue and tumours, suggests incremental increases in MYC activity at key transitions during early lung tumour emergence. Importantly, this is true even after removal of tumour samples expressing the Rosa26^DM.lsl-MYC^ allele from our analysis, indicating that endogenously expressed Myc, downstream of KRas^G12D^ activation, contributes substantially to the earliest stages of lung tumour development, and is consistent with data showing that loss of endogenous Myc suppresses KRas-driven lung tumourigenesis (68). The result is nonetheless surprising, given that activation of the Rosa26^DM.lsl-MYC^ allele in the absence of KRas^G12D^ expression fails to drive an overt neoplastic lung phenotype, even in the context of *Tp53* deletion (63). Modestly deregulated Myc expression is thus necessary but alone insufficient to initiate lung tumours in adult mice, contrasting with the proficiency of the same allele to alone drive neuroendocrine tumours when activated in the embryonic murine pancreas (11). A key difference between these models is that endogenously expressed Myc levels are high in the developing embryo but low in the adult lung (28), whereupon KRas activation in lung elevates Myc activity, either through protein stabilisation, increased gene expression, increased mRNA translation, or a combination of all three. Collectively, these data suggest that distinct threshold levels of MYC are required, first for neoplastic initiation in the lung, and subsequently to licence progression to frank tumour formation. Interestingly, the “high MYC” PDS1 subtype is not observed in KM mice carrying a single copy of the Rosa26^DM.lsl-MYC^ transgene but is in KM^2^ mice bearing 2 copies of the allele, indicating that a higher threshold level of MYC deregulation is also required to drive this phenotype. Accordingly, our analysis of the TCGA LuAd cohort shows MYC activity directly correlating with increasingly aggressive lung cancer, suggesting that MYC activity continues to rise incrementally through late LuAd progression, with a significant impact on patient survival.

All of which argues for the importance of developing a range of therapeutic (and arguably prophylactic) agents to counteract MYC activity. Several promising strategies are under active investigation, including small molecule and peptide disrupters of MYC:MAX binding, PROTAC degraders, Ubiquitin E3 ligase and Deubiquitylase inhibitors, and regulators of MYC transcription or translation, recently reviewed in excellent detail (29). Here we tested an orthogonal approach, using a small molecule targeting MYC-interacting cofactors, G9A and EZH2(62). Our rationale for doing so was largely based on the contribution of these cofactors to MYC-dependent transcriptional repression, now well-established to drive immune evasion through suppression of Type I Interferon signalling and MHC I antigen presentation across multiple cancer types (11, 18, 49, 69). Accordingly, HKMTi-1-005 treatment of autochthonous LuAd *in vivo* strongly increased expression of Type I Interferon regulators IRF7, IRF9 and STAT2, with more modest upregulation of MHC I related gene expression. This was accompanied by a pronounced increase in B cell-related gene expression, including gamma Immunoglobulins, indicative of B cell maturation. The precise role of B cells in lung tumour restraint is unclear but may involve cellular opsonisation and antibody-driven cytotoxicity, mediated by NK cells or other immune effectors. However, we found little evidence of T cell engagement following treatment with HKMTi-1-005, even when combined with PD1 blockade (Figure 4 & bioRxiv 2023.07.24.550274). Further work in immunogenic mouse models of autochthonous tumours will be needed to better understand the dynamics of T cell mediated anti-tumour activity in lung cancer.

Whole transcriptome analysis of autochthonous lung tumours additionally exposed a pronounced effect of HKMTi-1-005 on cell-cycle related gene expression, affecting both S phase and G2/M phase Cyclins.

MYC has previously been shown to repress expression of CDK inhibitors, p21 (CIP1) and p15 (INK4B), neither of which were significantly regulated in our analysis. Rather, this may suggest activity of the drug towards MYC transactivating complexes and indeed, EZH2 was previously shown to stabilise MYC protein (70). Accordingly, IHC analysis revealed strongly reduced MYC protein expression following treatment of lung tumours with HKMTi-1-005 (Fig. 5C), providing a likely explanation for the observed reduction in tumour cell proliferation *in vivo*. We recognise that both EZH2 and G9A have other roles in addition to their participation in MYC transcriptional regulatory complexes that may contribute to the cytostatic activity of the drug. Notably, EZH2 has been shown to participate in both RB-mediated gene silencing and E2F-dependent transactivation (71, 72), both intimately linked to Cyclin gene expression and cell cycle progression. However, EZH2 was also shown to prevent FBW7-mediated polyubiquitylation and degradation of MYCN and depletion of EZH2 accelerates MYCN turnover (70), and may function similarly here to regulate c-MYC stability. Importantly, sustained treatment with HKMTi-1-005 profoundly suppressed the growth of autochthonous lung tumours driven by elevated MYC expression and the cytostatic impact of drug was more pronounced in the model with higher MYC expression (i.e., KM^2^ versus KMA), in agreement with our *in silico* analysis of sensitivity to HKMTi-1-005 correlating with elevated MYC activity in human cancer cell lines.

We conclude that MYC transcriptional activity is a useful surrogate for stratifying LuAd patients, irrespective of driver mutation status, and moreover better defines early transition states of nascent neoplastic disease than absolute levels of MYC. We demonstrate that targeted inhibition of MYC-associated co-factors G9A & EZH2 can reduce MYC protein levels, suppress growth of MYC-driven LuAd and engage a measure of immune surveillance through upregulation of Type I Interferon signalling. This work thus establishes the principle of orthogonally targeting MYC, through inhibition of enzymatic co-factors, for treatment of LuAd.

## MATERIALS AND METHODS

### Mice and *In Vivo* Procedures

All mouse experiments were approved by the local Animal Welfare Ethics Review Board (AWERB) and conducted in accordance with UK Home Office licences PE47BC0BF and PP7768309 to DJM and Establishment licence XC2FD842E, to University of Glasgow. The *LSL-KRAS^G12D^*(53)*, Rosa26^DM.lsl-MYC^* (10)*, Hprt-lsl-IRFP* (73)*, Tp53^fl^* (54)*, Stk11^tm1.1Sjm^* (74) and *Rosa26^DS.lsl-APOBEC3B^* (bioRxiv 2023.07.24.550274) alleles were previously described. All mice were maintained on a mixed FVBN/C57Bl6 background, housed on a 12-hour light/dark cycle and fed and watered ad libitum. Male and female mice were included in all analyses in approximately equal numbers. Recombinant adenovirus expressing Ad5mSPC-CRE (SPC-CRE) was purchased from the University of Iowa gene therapy facility. To initiate lung tumours, 8–10-week-old mice were sedated with ketamine and medetomidine mixture via IP injection, followed by intranasal inhalation of SPC-CRE. For most experiments, 1x10^8^ viral pfu SPC-CRE were administered intranasally using the calcium phosphatase precipitation method, as described previously (9). For defined duration experiments, mice were aged to early or intermediate timepoints in tumour development. For survival analysis, cohorts of mice were monitored by facility personnel with no knowledge of experimental design and euthanised when humane endpoint was reached. All mice were humanely euthanised by CO_2_ inhalation followed by cervical dislocation. For histologic analysis, mouse lungs were perfusion-fixed in 10% neutral buffered formalin overnight. Prior to fixation, a small portion of tumour-bearing lung (approximately 30µg) was snap frozen for RNA from a subset of mice. For intervention analyses, mice were randomly assigned at time of tumour initiation to treatment or control groups, balanced only for sex. Drug administrator/investigator blinding was not performed for reasons of animal welfare. A dose titration of HKMTi-01-005 was performed on KM^2^ mice, in which mice were dosed twice daily with either 20mg/kg, 30mg/kg, 40mg/kg HKMTi-01-005, or vehicle control, for 5 consecutive days immediately prior to reaching 8-weeks post induction, then euthanised. To determine the impact of HKMTi-01-005 on tumour burden and proliferation in the KM^2^ model, mice were treated with 35mg/kg HKMTi-01-005 or vehicle control (0.9% NaCl) twice daily by IP injection on a schedule of 3 days on/4 days off from 4 weeks post induction to 8 weeks post induction and euthanised. To measure the potential survival benefit of HKMTi-01-005 in the KM^2^ model, mice were treated with 35mg/kg HKMTi-01-005 or vehicle control twice daily by IP injection on a schedule of 3 days on/four days off from 4 weeks post induction to 8 weeks post induction and euthanised at clinical endpoint. To determine the impact of HKMTi-01-005 on tumour infiltration by T cells, KMA mice were treated with 40mg/kg HKMTi-01-005, or vehicle control, twice daily by IP injection for 5 consecutive days immediately prior to 12 weeks post induction, then euthanised.

### Immunohistochemistry and Tissue Analysis

Mouse tissue IHC staining was performed on 4µm formalin-fixed, paraffin-embedded sections, which had been previously heated to 60°C for 2 hours. The following antibodies were stained on a Leica Bond Rx autostainer; CD45R (14-0452-82, Thermo Scientific), Rabbit IgG Isotype control (ab172730, Abcam) and Ki67 (12202, Cell Signalling Technology). All FFPE sections underwent on-board dewaxing (AR9222, Lecia) and epitope retrieval using epitope retrieval 2 solution (AR9640, Leica) for 20 minutes at 100°C. Sections were rinsed with Lecia wash buffer (AR9590, Leica) before peroxidase block was performed using an intense R kit (DS9263, Leica) for 5 minutes. Sections were rinsed with wash buffer before sections for CD45R had blocking solution applied from the Rat ImmPRESS kit (MP7444-15, Vector Labs) for 20 minutes. Sections were rinsed with wash buffer before primary antibody application at an optimal dilution (CD45R, 1/3000; IgG, 1/21000; Ki67, 1/1000). The sections were rinsed with wash buffer and sections being stained for IgG and Ki67 had Rabbit envision (K4003, Agilent) applied and sections for CD45R staining had Rat ImmPRESS secondary antibody applied for 30 minutes. The sections were rinsed with wash buffer, visualised using DAB and counterstained with haematoxylin in the Intense R kit. The IHC sections were rinsed in tap water, dehydrated through a series of graded alcohols and placed in xylene. The stained sections were coverslipped in xylene using DPX mountant (SEA-1300-00A, CellPath). Tumour area was calculated using HALO software (Indica Labs, v3.6.4134.396) as the percent area of the lung occupied by adenocarcinoma, measured on hematoxylin and eosin-stained sections. Ki67 positive cells were quantified using HALO software (Indica labs v3.6.4134.396). To stain for c-MYC expression, formalin fixed, paraffin embedded (FFPE) samples were sectioned using an automated microtome at 4µm, sections were placed onto TOMO adhesive microscope slides (Matsunami, 0808228600). Automated staining was performed on the Ventana Discovery Ultra platform (Roche Tissue Diagnostics RUO Discovery Universal V21.00.0019). Slides were baked at 60°C before a dewax step for 24 minutes at 95°C. Antigen retrieval (CC1, Roche Tissue Diagnostics, 06414575001) PH9 was performed for 32 minutes at 95°C. DISCOVERY Inhibitor CM (Roche Tissue Diagnostics, 05266645001) was applied for 12 minutes to block any endogenous peroxidase. c-MYC (Y69) (Roche Tissue Diagnostic 06504612001) Ready to Use (RTU) was applied for four hours at room temperature (RT). Discovery anti-Rabbit HQ (Roche Tissue Diagnostics, 07017812001) was applied for one hour followed by Discovery anti-HQ HRP (Roche Tissue Diagnostics 07017936001) for 32 minutes. DAB (Roche Tissue Diagnostics, 05266645001) was applied followed by Hematoxylin II (Roche Tissue Diagnostics, 05277965001) for 8 minutes and bluing reagent (Roche Tissue Diagnostics, 05266769001), for 4 mins. Slides were removed from the Ventana Discovery Ultra and washed in three rounds of Discovery wash (RUO 07311079001) followed by running tap water for one minute. Slides were transferred onto the Leica ST5010 Autostainer to be mounted with DPX (phthalate free) mounting medium (Cellpath SEA-1300-00A) and coverslipped (Solmedia COV52450).

### Bulk Tumour RNA Sequencing

To determine the transcriptomic impact of HKMTi-1-005 treatment on lung tumours, RNA was isolated from flash frozen KMA lung tumour samples using Qiagen RNeasy Kit according to the manufacturer’s protocol for purification of total RNA from animal tissues. DNA was depleted with the RNase-free DNase Set (Qiagen). The quality of purified RNA was tested on an Aligent 2200 Tapestation using RNA screentape. Libraries for cluster generation and DNA sequencing were prepared following an adapted method from (75), using Illumina TruSeq Stranded mRNA LT Kit. Quality and quantity of the DNA libraries were assessed on an Aligent 2200 Tapestation (D1000 screentape) and Qubit (Thermo Fisher Scientific) respectively. The libraries were run on the Illumina Next Seq 500 using the High Output 75 cycles kit (2 x 36 cycles, paired end reads, single index). Quality checks and trimming on the raw RNA-Seq data files were done using FastQC version 0.11.9 (https://www.bioinformatics.babraham.ac.uk/projects/fastqc/), FastP version 0.20.1 (76), and FastQ Screen version 0.14 (77). RNA-Seq paired-end reads were aligned to the GRCm39.105 version of the mouse genome and annotation (78), using HiSat2 version 2.2.1 (79) and sorted using Samtools version 1.7 (80). Aligned genes were identified using Feature Counts from the SubRead package version 2.0.1 (81). Expression levels were determined and statistically analysed using the R environment version 4.1.2 (82) and utilising packages from the Bioconductor data analysis suite (83). Differential gene expression was analysed based on the negative binomial distribution using the DESeq2 package version 1.34.0 (84) and adaptive shrinkage using Ashr (85). Identification of enriched biological functions was achieved using g:Profiler (86), and GSEA version 7.4 from the Broad Institute (87). Computational analysis was documented at each stage using MultiQC v1.11 (88), Jupyter Notebooks and R Notebooks (89).

For cross-genotype comparison of tumour phenotypes, we generated a tissue microarray (TMA) of visually selected tumours from the lungs of end-stage KM, KM^2^, KMA, KL, KLA, KP and KPA mice (see Figure S2). The TMA comprised 4 blocks of “cored” tissue samples and included a small number of liver metastases along with adjacent liver samples that were excluded from the analysis performed here. Formalin fixed paraffin embedded tissue sections (4μm thick) were mounted on positively charged glass microscopic slides and air dried at room temperature overnight. The sections were incubated for 30 minutes in a 60°C oven immediately prior to undergoing heat mediated epitope retrieval (Leica BOND RX; HEIR 10 minutes with ER2 at 100°C) and 1µg/ml proteinase K (AM2546, Thermo Fisher Scientific) digestion for 15 minutes at 37°C. The sections were incubated overnight in a hybridisation oven at 37°C with RNA oligo detection probes (GeoMx Mouse Whole Transcriptome Atlas, Nanostring, targeting over 21,000 transcripts for mouse protein coding genes and negative controls). Morphological structures were stained using a fluorescent labelled pan cytokeratin antibody (1:1000, NBP2-33200-AF532, Novus) and a fluorescent nucleic acid stain (1:10,000, SYTO13, Nanostring). Slides were imaged at 20x magnification using the GeoMx Digital Spatial Profiler (DSP). Through the integrated DSP software suite, 232 regions of interest (ROIs) were selected (see Table S1) and UV illuminated (385 nm), cleaving the UV-sensitive linker in the RNA detection probes and allowing for subsequent collection of the oligo barcode component in a 96-well plate. Libraries were prepared using GeoMx Seq Code primers (NanoString) and 1x PCR Master Mix (NanoString). Samples from each collection plate were equivolume-pooled into a plate-corresponding sample pool and AMPure XP purified. Library quality was assessed using an Agilent Bioanalyser. The libraries were sequenced on an Illumina NovaSeq 6000 System (GeneWiz/Azenta) at the assay recommended sequencing depth (number of sequenced read pairs = total collection area (mm^2^) x Sequencing Depth Factor [100 for Whole Transcriptome Atlas]). Raw sequencing reads were processed using the GeoMx NGS Pipeline (v2.3.3.10) to generate Digital Count Conversion (DCC) files. Subsequent data management and formatting were conducted in the R statistical environment (v4.5.1). Project configuration and file path management were handled using the yaml package (v2.3.10). Experimental metadata, including segment properties, probe count matrices, and laboratory worksheets, were imported, standardised and merged using the readxl (v1.4.5) and tidyverse (v2.0.0) packages to construct a unified dataset for downstream analysis. Data loading and quality control (QC) were performed using the Geomx Tools package (v3.6.2) and NanoStringNCTools (v1.10.0). Raw DCC files were collated and merged with PKC assay definition files and sample annotations. Segment QC was assessed based on sequencing saturation, raw read counts, and negative control background levels. Probes were filtered to remove low-performing targets falling below the limit of quantification (LOQ), defined as the geometric mean of negative control probes multiplied by a geometric standard deviation factor (typically 2.5). To account for technical variation and batch effects arising from the TMA structure, normalisation was conducted using the standR package (v1.0.0). The Remove Unwanted Variation (RUV algorithm) was applied to correct for technical artefacts while preserving biological heterogeneity across genotypes and tissue types. Visualisation of QC metrics and principal component analysis (PCA) were performed using ggplot2 (v3.5.2) and scater (v1.30.1) to assess data structure post-normalisation. ROIs from 1 of the 4 slides exhibited significant divergence from the rest of the sample set and the entire slide (identified as TMA2) was excluded from further analysis. One further ROI was excluded on the grounds of extreme transcriptional divergence, likely owing to unintentional inclusion of a suspected tertiary lymphoid structure. Differential expression analysis was conducted using the DESeq2 package (v1.42.1), utilising a negative binomial generalised linear model to compare transcriptomic profiles across phenotypes (Tumours, Hyperplastic, Normal Lung) and genotypes. To assess significance, P-values were adjusted for multiple testing using the Benjamini-Hochberg procedure (FDR < 0.05). Wald confidence intervals were calculated for log2 fold changes to provide robust effect size estimates. Gene Set Enrichment Analysis (GSEA) was performed using the fgsea package (v1.28.0) to identify enriched biological pathways from the MSigDB Hallmark gene sets. Additionally, single-sample GSEA (ssGSEA) was applied using the escape package (v1.12.0) to derive pathway activity scores from individual ROIs, enabling sample-level comparisons. Results were visualised using a suite of R packages. Volcano plots and MA plots were generated using ggplots2 and ggrepel (v0.9.5) to display differential expression results. Complex heatmaps of gene expression and pathway activity scores were constructed using ComplexHeatmap (v2.18.0) and pheatmap (v1.0.12), employing hierarchical clustering to reveal patterns across sample groups. Dimensionality reduction (PCA, t-SNE, UMAP) plots were created to visualise sample separation based on transcriptome profiles.

### Cell lines and cell culture

A549 and H358 cells were obtained from ATCC and cultured in RPMI 1640 containing 10% FCS, 2mM L-glutamine (Thermo Fisher Scientific) and 10,000 U/ml penicillin/streptomycin (Thermo Fisher Scientific). Cells in log phase growth were treated with indicated concentrations of HKMTi-01-005, UNC1999 (HY-15646; MedChem Express), UNC0642 (HY-13980; MedChem Express) or MYCi975 (S8906; Selleckchem) throughout the study. Equivalent volumes of diluents were used as vehicle controls. For immunoblotting, whole cell lysates were prepared in RIPA buffer (150mmol/L NaCl, 50mmol/L Tris pH7.5, 1% NP-40, 0.5% sodium deoxycholic acid, 1% SDS, plus complete protease and phosphatase inhibitor cocktail) followed by sonication (40% Amp for 10 seconds). MYC (ab32072) and Vinculin (ab129002) were used as primary antibodies. Secondary HRP-conjugated antibodies (α-Mouse IgG NA931V and α-Rabbit IgG NA934V, both GE Healthcare) were detected by chemiluminescence (Bio-Rad Western Blotting substrate 1705060). To measure confluency, an IncuCyte (Sartorious, Gottingen, Germany) was used to monitor growth over time. A549s and H358s were plated at a seeding density of 5,000 and 10,000 cells per well, respectively, in a 96-well plate. The following day, drugs were added in fresh media and proliferation was monitored for 72 hours. To detect interferon activator gene expression, cDNA was synthesized with the QuantiTect Reverse Transcription Kit (Qiagen) and pre-amplified (20nmol/L of each primer and SYBR green buffer) for *GUSB* (F cctgtgacctttgtgagcaa; R aacagatcacatccacatacgg), *IRF7* (F ctgcagtcacacctgtagcc; R gtggactgagggcttgtagc), *IRF9* (tcctccagagccagactact; R caatccaggctttgcacctg), *STAT1* (F ctccactgccactgcgctcc; R gccccttggcactcctgctg) and *STAT2* (F accattctggacatggctgg; R ctccgactcacaaagcccat) for 20 cycles. Diluted cDNA was quantified by real time PCR using SYBR Green method (VWR QUNT95072).

### Statistical Analysis

Raw data obtained from qRT-PCR and growth curves were copied into GraphPad Prism, where graphical representation of the data was produced and all mean and SEM calculations were calculated. Statistical significance was determined by the Student’s T test. For multiple comparisons, ANOVA was used with a post hoc Tukey test. For non-normally distributed data (e.g. survival data), a Mantel-Cox (two-way comparison) test was performed. For RNA-seq data, adjusted P values calculated in R Studio are shown. For all panels, * denotes P <0.05; **, P <0.01; ***, P < 0.005; ****, P < 0.001.

### Computational Biology – TCGA LUAD analysis

Transcriptional and mutational data for primary tumour samples from The Cancer Genome Atlas lung adenocarcinoma (TCGA LUAD) cohort were accessed from the Genomic Data Commons (GDC) via the TCGAbiolinks (90) R package on 21st July 2022. RNA-seq data was processed using DESeq2 (84), with variance stabilising transformation (VST) performed through the vst function. Mutational status for genes, in each of the TCGA LUAD samples, was determined according to the MAF file downloaded through TCGAbiolinks. Samples harbouring non-silent mutations in a given gene were classified as mutant; others were classified as wild type (WT). Samples were classified according to the PDS subtyping system by applying the PDSpredict function from the PDSclassifier package (51), to the VST-transformed RNA-seq data, using the default classification threshold of 0.6. Hallmark gene signatures were accessed from Molecular Signatures Database (MSigDB) (52) using msigdbr package (91). Single sample gene set enrichment analysis (ssGSEA), as implemented by the GSVA package (92), was performed on the VST-transformed RNA-seq data to calculate enrichment scores for the 50 Hallmark gene sets. Proliferative index was calculated using the ProliferativeIndex R package (93). Replication stress was assessed using n=20 transcriptional signatures associated with cell cycle and DNA repair, as previously described (51, 94). These signatures were obtained from MSigDB4 via the msigdbr package5. Enrichment scores for these signatures were calculated using the gsva method, from the GSVA R package (92), on the VST-transformed RNA-seq data. For each sample, the sum of GSVA enrichment scores for the 20 signatures was used as the replication stress score. For transcription factor activity inference, the DoRothEA (95) collection of transcription factor-target interactions (regulons) was used, specifically the high confidence (A and B) regulons. The run_viper function from the dorothea R package was applied to the VST-transformed RNA-seq data to infer transcription factor activity scores. Overall survival analysis was performed based on the vital status, days to death and days to last follow up variables, in the clinical data obtained for TCGA LUAD through TCGAbiolinks, with censoring at five years. Samples missing the required survival data (n=9) were excluded from these analyses. Differences in survival between groups were tested for using log-rank tests, through the survminer package.

### Computational Biology – Sanger Cell Line analysis

Drug sensitivity data in the form of IC_50_ values were downloaded from the Genomics of Drug Sensitivity in Cancer 2 (GDSC2) resource (https://www.cancerrxgene.org/downloads/bulk_download, accessed 30/09/22). Methods for calculating IC_50_ values based on cell viability assays are described in the accompanying documentation available at the same site. Raw data downloaded included sensitivity measurements for 295 compounds, across 969 cell lines derived from many different tumour types. Within these data, there were two single G9a (EHMT2) inhibitors (UNC0638 and A-366, DRUG IDs 2038 and 2157 respectively), and two single EZH2i studies (GSK343 in two separate screens, DRUG IDs 1627 and 2037), alongside HKMTi-1-005. Of these, we selected UNC0638 for a G9ai comparison, as an inhibitor with the same substrate-competitive mechanism of action as HKMTi-1-005. For EZH2i comparison, we included the GSK343 data with the greatest overlap in cell lines tested as compared to HKMTi-1-005 (DRUG ID 2037). Cell lines from solid tumours with sensitivity data for all four compounds of interest were included in analyses (n = 565). Gene expression (in the form of RMA data), mutation status and amplification data were downloaded from the COSMIC Cell Lines Project v96, released 31/05/22. Copy number alteration such as those recorded for c-MYC are defined by COSMIC as an amplification if the average genome ploidy <= 2.7 and total copy number >= 5, or the average genome ploidy > 2.7 and the total copy number >= 9. Genesets included in our analysis are listed in Table S2. Interferon Alpha Response, Interferon Gamma Response and Innate Immune Response genesets were downloaded from MSigDB, (https://www.gsea-msigdb.org/gsea/msigdb/human/genesets.jsp v7.4), accessed 20/10/22 and 22/03/23. The cMYC-G9a repression geneset was created based on literature published describing a list of 38 genes repressed by this complex in breast cancer cell lines (41, 96). The dual EZH2/G9a target geneset was defined by Casciello et al. (97). The singscore package was used to score samples for these genesets (98, 99) based on RPKM (ICGC), TPM (TCGA) or RMA counts (GDSC2) to ensure correction for gene length bias. RPKM/TPM counts were filtered prior to scoring, to ensure > 0.5 RPKM/TPM in more than half the samples. *Statistical analysis:* Unless otherwise stated, graphs and statistics were generated in R (82) and RStudio version 4.2.2 (89). Key packages besides those described below included ggplot2 (100) and tidyverse (101). If the data were normally distributed, or could be transformed to be so, correlation analyses were performed using a Pearson’s analysis. If non-normal, correlations were performed using a Spearman’s analysis. Groupwise comparisons were performed using t-tests or Mann-Whitney/Wilcoxon tests, for normal or non-normal data respectively, using ggpubr and ggstatsplot packages (100, 102). Multiple linear regression was performed using the lm function from the R base package stats (82), with added-variable plots for visualisation generated using the car package (103). Lung cancer cell line dependency data following CRISPR-mediated *MYC* deletion were downloaded from the Depmap Achilles Project portal version 25Q2 (https://depmap.org/portal/achilles/) and visualised using Graphpad Prism.

## Data & Coding Availability

FASTQ files and related metadata pertaining to GeoMX whole transcriptome analysis of mouse lung tumour tissue microarrays are available from ArrayExpress, project E-MTAB-16376. FASTQ files and related metadata pertaining to HKMTi-1-005 treatment of KMA mice are available from ArrayExpress, project E-MTAB-16350. Genesets used in the analysis of GDSC2 cell lines are listed in Table S2. Code related to whole transcriptome analysis of mouse lung tumours is available at https://github.com/rwsBiomics/geomx-lung-myc-gemm.

## Material Availability

The Rosa26^DM.lsl-MYC^ allelic mice are available from Jaxmice, RRID:IMSR_JAX:033805. The Rosa26^DS.lsl-APOBEC3B^ allelic mice are available upon request from the CRUK SI.

## CREDIT Contributions

**Conceptualization –** SL, ID, PD, RB(s) & DJM

**Data curation –** SL, ID, RB(j), EF, RS & SA

**Formal analysis –** SL, ID, RB(j), EF, RS, & SA

**Funding acquisition –** DJM, RB(s), PD, NBJ, JLQ & KB

**Investigation –** SL, ID, RB & SH

**Methodology –** SL, ID, RB(j), EF, RS, SA, YD, CF, JD, SMcL, CN, GC, LOJ, CKD & NB

**Project administration –** SL, PD, RB(s) & DJM

**Resources –** SBM, SG, SMcL, DS, IMcN & MJF

**Software –** RS, SA, PD

**Supervision –** IMcN, CM, KB, JLQ, NJ, PD, RB & DJM

**Validation –** SL, ID, RS, EH, PD

**Visualization –** SL, ID, RB(j), RS, SA & DJM

**Writing – original draft –** SL, ID, RB(j), PD, RB(s) & DJM

**Writing – reviewing & editing –** All authors

## ACKNOWLEDGEMENTS

The authors wish to thank the staff of the CRUK SI Biological Services Unit for all animal husbandry, routine monitoring and procedural assistance. We similarly thank the staff of the CRUK SI Histology Core Facility and the Deep Phenotyping Advanced Technology Core Facility (RRID:SCR_027366). Special thanks to CRUK SI Integrity Officer, Dr. Catherine Winchester, for careful evaluation of the manuscript and guidance on best publishing practice.

## Associated Pre-prints

Data included in this manuscript were previously uploaded to Biorxiv as part of 2 separate manuscripts: DOI: 10.1101/2023.10.18.562888 (Figure 3, panels C-E; Figure 5, panels E-H; Figure S4, panels A, B) DOI: 10.1101/2023.07.24.550274 (Figure 4, panels D-H; Figure S5, panels A-C)

## SUPPLEMENTARY FIGURES

**Figure S1.**
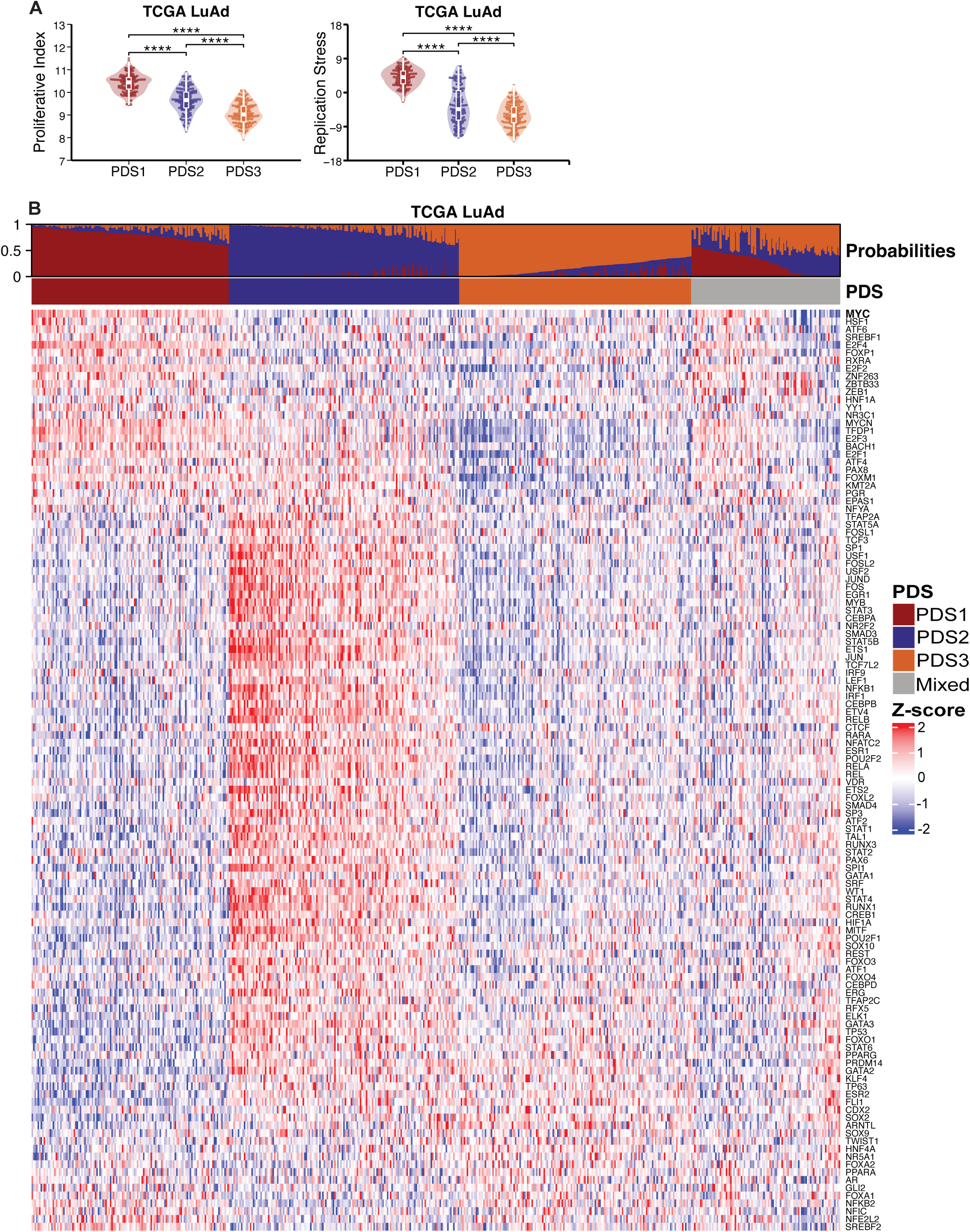
**Relates to Main Figure 1** **A)** Proliferative index (left) and replication stress scores (right) according to PDS in TCGA LuAd. Each dot represents one tumour sample. Boxplots indicate interquartile range (25th–75th percentiles) with median values represented by the central horizontal line. Whiskers extend to values within 1.5 times the interquartile range of the upper and lower quartiles. Observations beyond the end of the whiskers are plotted individually. Mann-Whitney test values indicated. **B)** Heatmap showing inferred transcription factor activity (TFA) scores in the TCGA LuAd samples, with samples arranged according to PDS. Annotation bars above heatmap indicating the PDS subtype of samples and associated predicted PDS probabilities. Mann-Whitney U test values indicated. For all figures, **** = p < 0.0001, *** = p < 0.001, ** = p < 0.01, * = p < 0.05, and ns = p > 0.05.

**Figure S2.**
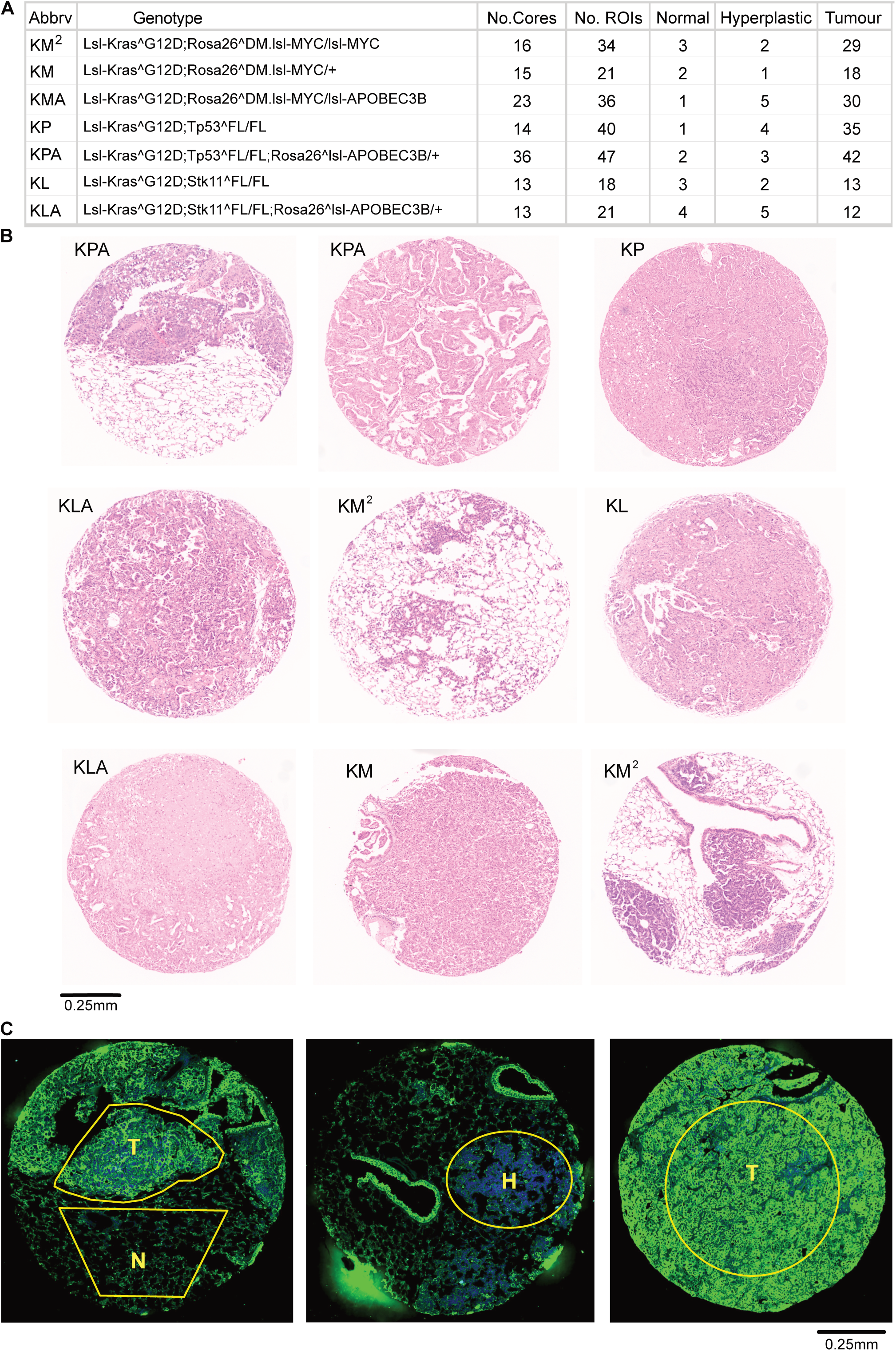
**Relates to Main Figure 2** **A)** Table indicating number of cores, regions of interest (ROIs) and tissue type selected for analysis in GeoMx Digital Spatial Profiler pipeline. **B)** Selected images showing varied histology of lung tumours from end-stage mice of various genotypes. Images are not intended to be representative of each genotype and end-stage tumours from each genotype show a multitude of histological appearances. **C)** Representative immunofluorescent images of lung tumours stained with pan-Cytokeratin antibody (Green) and Syto13 (Blue), showing examples of tumour (T), hyperplastic (H) and adjacent normal (N) tissue harvested for whole transcriptome analysis.

**Figure S3.**
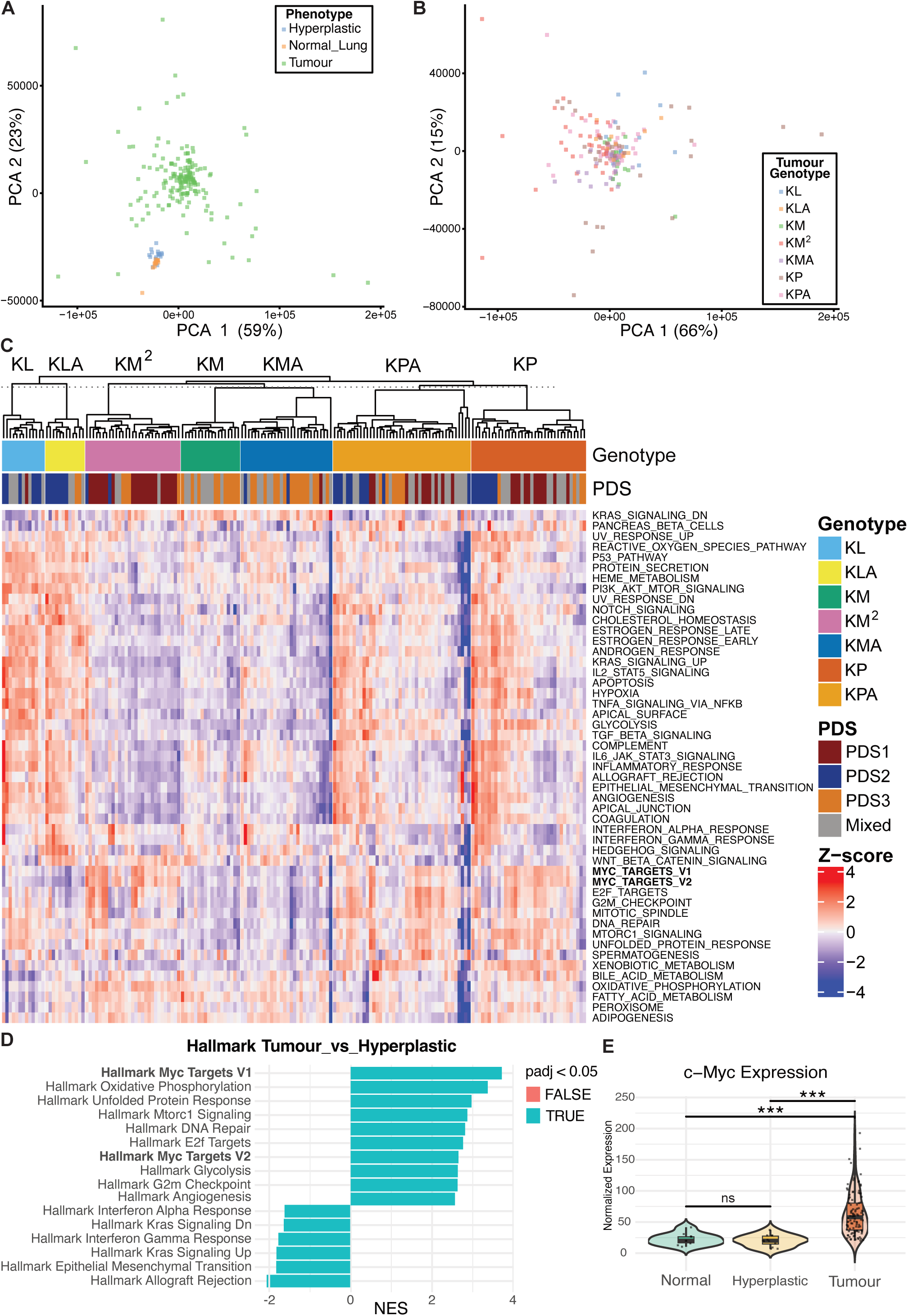
**Relates to Main Figure 2** **A)** PCA plot showing the clustering of tissue phenotypes from mouse lung TMAs. N=16 normal tissue, 22 hyperplastic tissue, and 179 tumour regions of interest (ROIs) selected for sequencing analysis. **B)** PCA plot showing the clustering of tumour genotypes selected for sequencing from mouse lung TMAs. N number of tumour ROIs for each genotype as shown in Figure S2A. **C)** Heatmap showing ssGSEA scores for all 50 Hallmark gene sets in tumour samples from TMAs of LUAD mouse models, from N=231 tumour ROIs organised by genotype of the mouse model. Annotation bars above the heatmap indicate the genotype of the mouse model and the PDS the tumours stratify to. **E)** Differential expression analysis between lung tumour tissue (N=126) and hyperplastic lung tissue (N=22) showing ssGSEA scores for Hallmark pathways that are differentially expressed in mouse ROIs excluding Rosa26^DM.lsl-MYC^ positive tumour samples. FDR<0.05 indicated by turquoise bars. **F)** Comparison of normalised endogenous *c-Myc* expression in normal, hyperplastic and tumours ROIs from lungs of mice, excluding those carrying the *Rosa26^DM.lsl-MYC^* allele. 1 way ANOVA with post-hoc Tukey test. *** denotes p<0.001.

**Figure S4.**
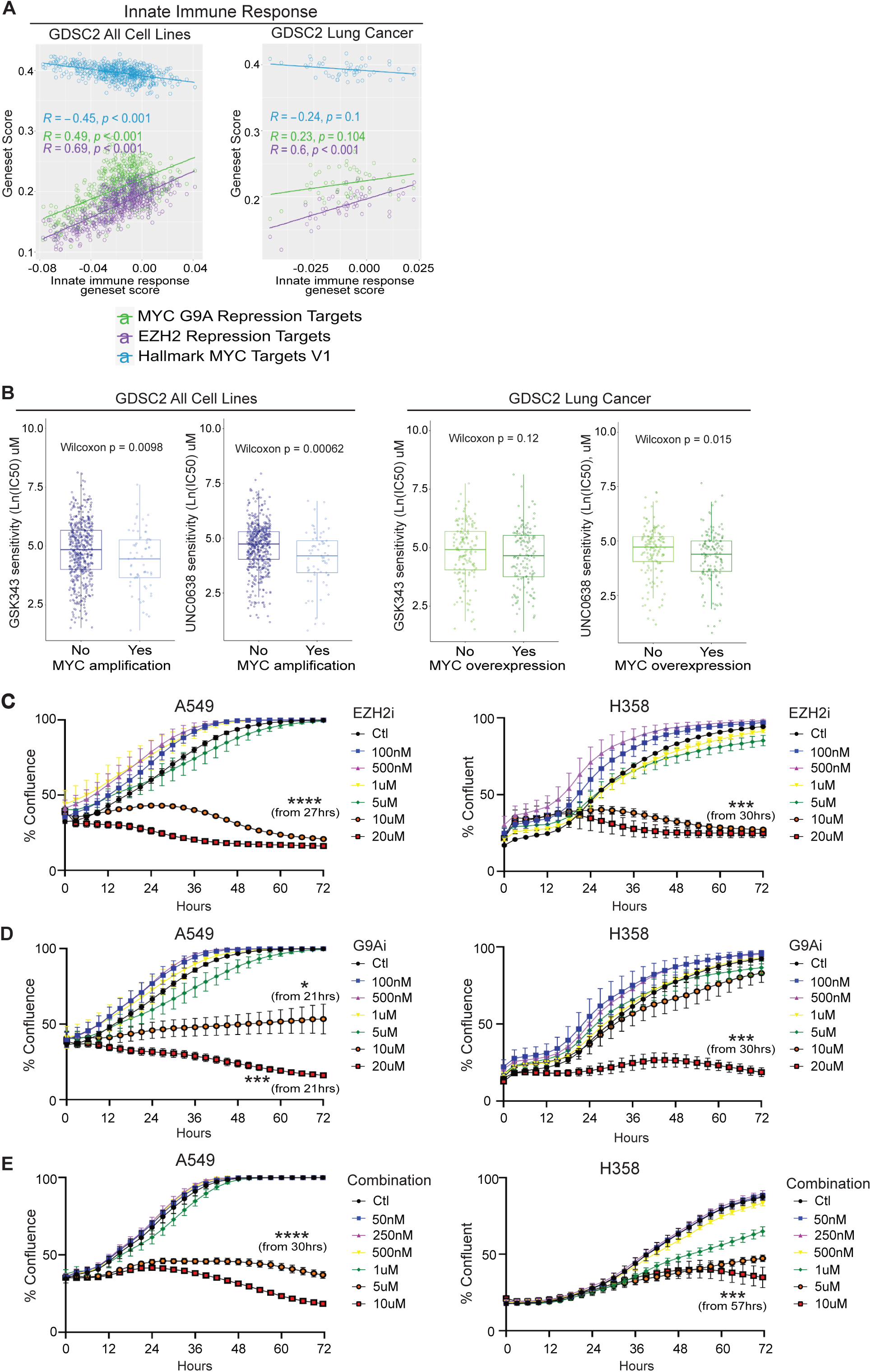
**Relates to Main Figure 4** **A)** Spearman’s correlation between Innate Immune Response geneset (x axis) and genesets reflecting targets of cMYC-G9a repression, EZH2 repression, and hallmark cMYC activation (y axis), in all GDSC2 solid tumour cell lines (left panel; N = 565) or GDSC2 lung cancer cell lines (right panel; N = 49). **B)** Difference in sensitivity to G9A inhibitor (UNC0638) or EZH2 inhibitor (GSK343) between *cMYC*-amplified (N = 65) and non-amplified (N = 500) solid tumour cell lines (left 2 panels) and between highest and lowest quartiles for MYC expression (right 2 panels; N = 141 each). Wilcoxon test. **C & D)** Confluence of A549 and H358 cells treated with 100nM, 500nm, 1µM, 5 µM, 10 µM & 20 µM GSK343 (EZH2i) **(C)** or UNC0638 (G9Ai) **(D)** or DMSO vehicle, measured over time by incucyte. Mean values ± SEM of 9 technical replicates (3 image fields x 3 replicates wells) from 1 experiment shown. Data are representative of four independent experiments. Significance calculated using a one-way ANOVA with post-hoc Tukey test for multiple comparisons. **E)** Confluence of A549 and H358 cells treated with a combination of EZH2i and G9Ai at individual concentrations of 50nM, 250nM, 500nM, 1µM, 5µM and 10µM, or DMSO vehicle measured over time by Incucyte. Mean values ± SEM of 9 technical replicates (3 image fields x 3 replicates wells) from 1 experiment shown. Data are representative of four independent experiments. Significance calculated using a one-way ANOVA with post-hoc Tukey test for multiple comparisons.

**Figure S5.**
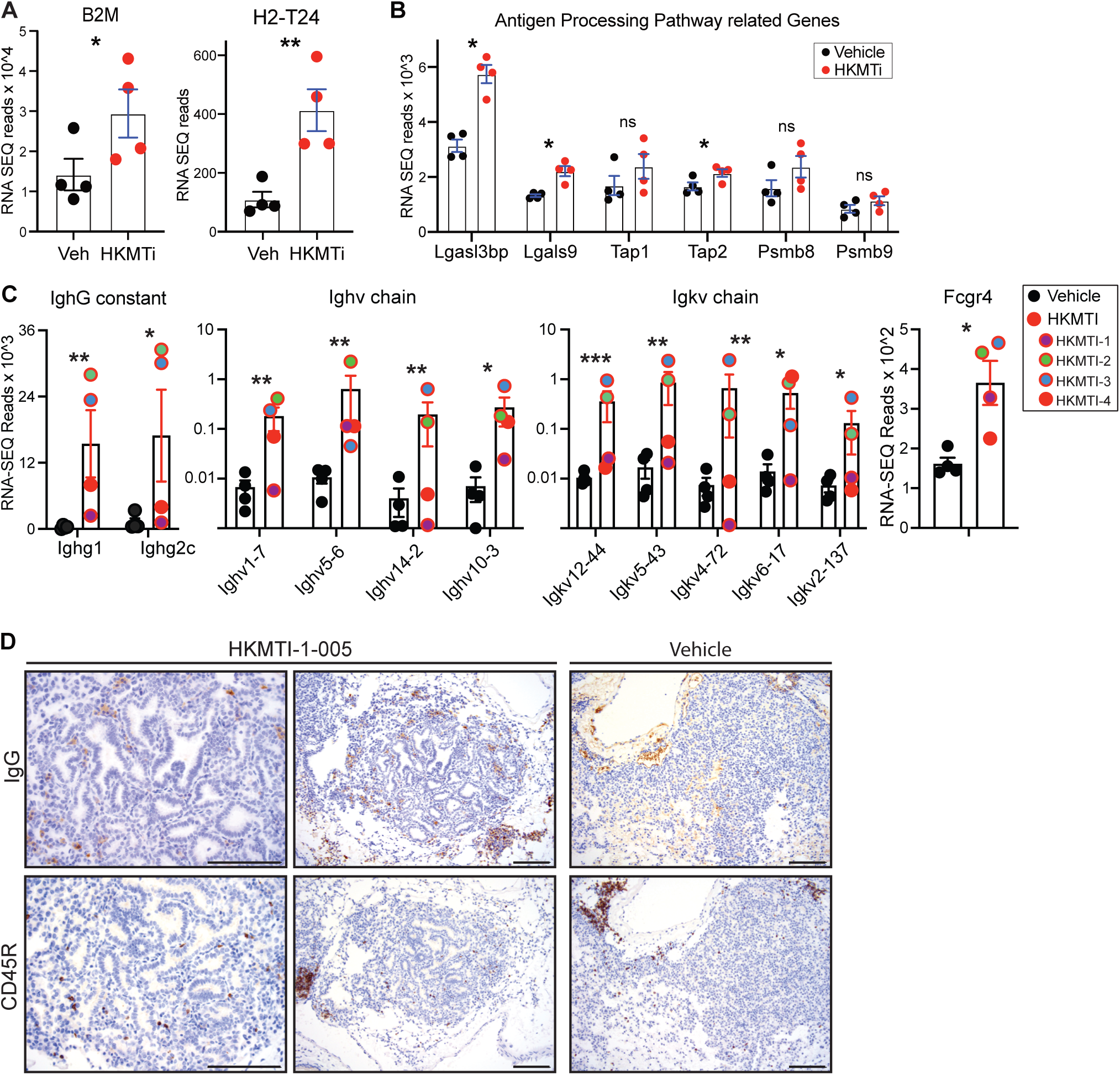
**Relates to Main Figure 4** **A+B)** Normalised RNA-Seq reads of MHC I antigen processing and presenting genes from tumour bearing lungs from KMA mice treated with vehicle (N = 4) or 40mg/kg HKMTI-1-005 (N = 4) for 5 days prior to cull at 12-weeks post allele induction. Mean and SEM shown. P values adjusted for multiple comparisons. **C)** Normalised RNA-Seq reads of IgG heavy and light chain genes from KMA mice as in S5A+B. **D)** Representative images of IgG and CD45R stained KMA lungs from mice as in S5A+B. Scale bar = 100µM.

**Table S1**

Metadata for all ROIs selected from GEMM end-stage lung TMAs (supplementary Excel file)

**Table S2**

Genesets used for cell line scoring of pathway correlations and drug sensitivity

**Table S2:** Genesets used for GDSC2 pathways analysish

| Geneset | Description | Source | Number of genes in geneset |
| --- | --- | --- | --- |
| (GOBP)INNATE_IMMUNE_RESPONSE | Innate immune responses are defense responses mediated by germline encoded components that directly recognize components of potential pathogens.<br>[GO_REF:0000022, GOC:add, GOC:ebc, GOC:mtg_sensu] | MSigDB v2023.1.Hs | 977 |
| EZH2_REPRESSION_TARGETS | Derived from a meta-analysis of 18 independent microarray studies performed by Curry et al., consensus genes that are upregulated by EZH2 KD in a variety of cell lines. | Curry et al., Clin Epigenetics. 2015 Aug 21;7(1):84. doi: 10.1186/s13148-015-0118-9. PMID: 26300989; PMCID: PMC4545913. | 300 |
| HALLMARK_INTERFERON_ALPHA_RESPONSE | Genes up-regulated in response to alpha interferon proteins. | MSigDB v2023.1.Hs | 97 |
| HALLMARK_INTERFERON_GAMMA_RESPONSE | Genes up-regulated in response to IFNG [GeneID=3458]. | MSigDB v2023.1.Hs | 200 |
| HALLMARK_MYC_TARGETS_V1 | A subgroup of genes regulated by MYC - version 1 (v1). | MSigDB v2022.1.Hs | 200 |
| MYC_G9A_REPRESSION_TARGETS | Defined by Tu et al., genes that had decreased cMYC binding according to ChIPseq whilst also being upregulated at the gene expression level by RNAseq analysis, in MCF10A.PM and MDAB231 cell treated with short hairpin targeting G9a. | Tu et al., Cancer Cell. 2018 Oct 8;34(4):579-595.e8. doi: 10.1016/j.ccell.2018.09.001. PMID: 30300580. | 38 |

